# The fitness consequences of genetic divergence between polymorphic gene arrangements

**DOI:** 10.1101/2023.10.16.562579

**Authors:** Brian Charlesworth

## Abstract

Inversions restrict recombination when heterozygous with standard arrangements, but often have few noticeable phenotypic effects. Nevertheless, there are several examples of inversions that can be maintained polymorphic by strong selection under laboratory conditions. A long-standing model for the source of such selection is divergence between arrangements with respect to recessive or partially recessive deleterious mutations, resulting in a selective advantage to heterokaryotypic individuals over homokaryotypes. This paper uses a combination of analytical and numerical methods to investigate this model, for the simple case of an autosomal inversion with multiple independent nucleotide sites subject to deleterious mutations. A complete lack of recombination in heterokaryotypes is assumed, as well as constancy of the frequency of the inversion over space and time. It is shown that a significantly higher mutational load will develop for the less frequent arrangement. A selective advantage to heterokaryotypes is only expected when the two alternative arrangements are nearly equal in frequency, so that their mutational loads are very similar in size. The effects of some *Drosophila pseudoobscura* polymorphic inversions on fitness traits seem to be too large to be explained by this process, although it may contribute to some of the observed effects. Several population genomic statistics can provide evidence for signatures of a reduced efficacy of selection associated with the rarer of two arrangements, but there is currently little published data that are relevant to the theoretical predictions.

## Introduction

Wright and Dobzhansky (1946) obtained evidence that some natural inversion polymorphisms in *Drosophila pseudoobscura* are associated with major differences in fitness among karyotypes, which can lead to their stable maintenance within a single population under constant environmental conditions. There have subsequently been many other experimental studies documenting strong effects of inversion karyotypes on fitness components in several *Drosophila* species (reviewd in Krimbas and Powell 1992; Kapun and Flatt 2019), and in some other species such as the seaweed fly *Coelopa frigida* (Butlin et al. 1984; Mérot et al. 2020). The startling observations of Wright and Dobzhansky (1946) raised the question of the causes of fitness differences between apparently functionally insignificant chromosomal variants. This question is still the subject of ongoing inquiry, stimulated by the new evidence from genome sequencing that inversion polymorphisms are abundant in natural populations of many species (Wellenreuther and Bernatchez 2018; Faria et al. 2019; Berdan et al. 2023).

Well before the work of Wright and Dobzhansky, Sturtevant and Mather (1938) had proposed a process that could result in fitness differences between inversion karyotypes, leading to a fitness advantage to heterokaryotypes over homokaryotypes. In their words: “… if a chromosome exists, in a population, in two sequences, differing by an inversion, it will in effect show two distinct lines of descent. There is free exchange of material within any line (*i*.*e*., sequence), but none between the sequences. Therefore, fluctuations of the genic contents must occur almost independently in the two sequences. Under such conditions, it is inevitable that in time the gene content of the two sequences will become different. It must be supposed that each sequence is susceptible to the same mutations, and with the same frequencies, but, as a result of the relative rarity in the population of a given mutant allelomorph at any one moment, certain genes will be present in one sequence but not the other. It is thus clear, considering two sequences A and B, that the homozygotes AA and BB are more likely to be homozygous for deleterious recessive mutations than is the heterozygote AB. Thus in general the sequence heterozygote AB will be at a selective advantage with respect to either of the corresponding homozygotes.”

Sturtevant and Mather (1938) did not attempt a quantitative model of this process, simply noting that “there can be no stability in the exact relations of the sequences with each other”, and that “…. a single gene difference can not in general cause such heterosis. The simplest effective condition is that in which each sequence contains a deleterious recessive not present in the other.” This proposal raises the question of what strength of selection on the two arrangements might be expected on its basis.

Ohta (1971) developed a mathematical model of a closely related, but not identical, process, based on the concept of associative overdominance (AOD), which was first outlined by Frydenberg (1963). Here, a polymorphic neutral locus can acquire an apparent heterozygote advantage, because of linkage disequilibrium (LD) generated by genetic drift with a locus subject to selection in favor of heterozygotes or to selection against recessive/partially recessive deleterious alleles maintained by mutation pressure. Unlike the model of Sturtevant and Mather (1938), this process does not require the generation of heterosis by multiple selected loci, which is often referred to as pseudo-overdominance (Waller 2021). Ohta (1971) summed the effects of individual selected loci that were completely linked to a dialellic neutral locus (equivalent to an inversion polymorphism) over a large segment of genome, and generated expressions for the apparent fitness advantage to heterozygotes at the neutral locus. This advantage could be substantial under suitable conditions with respect to population size, selection intensity, dominance and mutation rate.

It was, however, shown by Zhao and Charlesworth (2016) that Ohta’s formulae for the relative fitnesses at the neutral locus induced by linkage disequilibrium with the selected locus do not predict any change in allele frequency at the neutral locus, because they use only the squares of the linkage disequilibrium coefficient (*D*), whereas the allele frequency change involves the product of *D* and the additive effect on fitness of the selected locus (the Price equation: Price 1970). An induced selection pressure in favor of increased variability at the neutral locus only exists when the product of population size and selection coefficient is of the order of one; otherwise variability is reduced by background selection effects, even for recessive or partially recessive deleterious mutations (see also Charlesworth 2022). Ohta’s results therefore do not solve the quantitative problem of whether the strength of selection on inversions revealed by the experiments cited above can be explained by this process. Nonetheless, it is still often invoked as a potential contributor to the selective maintenance of inversion polymorphisms, e.g., Faria et al. (2019), Berdan et al. (2021), Jay et al. (2021) and Matschiner et al. (2023).

A somewhat different perspective was developed by Nei et al. (1967), who examined a purely deterministic model involving the balance between mutation and selection at numerous autosomal loci. This process results in an equilibrium frequency distribution of the number of mutant alleles per haploid genome; a new autosomal inversion has a reasonable chance of arising on a haplotype with a lower number of mutations than average; but, as shown by Nei et al. (1967), as time goes on, the mutant-free loci on the inverted haplotypes will accumulate mutations; unless the inversion goes to fixation, the loci in the inversion subpopulation will eventually acquire the same frequencies of mutant alleles as the corresponding loci in the subpopulation with the standard arrangement. Nei et al. (1967) interpreted this as implying that the inversion would then be selectively neutral. However, if reverse mutations from mutant to wild-type alleles do not occur, the loci at which mutations were present in the original inversion haplotype will all be homozygous in inversion homokaryotypes, causing a reduced fitness compared to that of homokaryotypes for the standard arrangement. A reanalysis and extension of this model by Connallon and Olito (2021) suggested that it can result in a net heterozygote advantage to an autosomal inversion in a randomly mating population, provided that inversions have a sufficiently large direct selective (but non-heterotic) advantage that is independent of the deleterious mutations, resulting in the maintenance of the inversion by selection.

Berdan et al. (2021) conducted simulations of a finite population with multiple loci experiencing mutations to deleterious and completely recessive alleles and found that a selective advantage to heterokaryotypes could develop, provided that recombinational exchange between arrangements in heterokaryotypes was sufficiently infrequent, and a small additional selective advantage to heterokaryotypes kept the inversion in the population long enough for mutation accumulation to occur. However, extensive computer simulations of a multi-locus model of mutation and selection with no selective bonus and less extreme assumptions about the degree of recessivity of deleterious mutations by Jay et al. (2022) showed that autosomal inversion polymorphisms are unlikely to be established. Studies of the effects of deleterious mutations have shown that complete recessivity is unlikely to be frequent, especially for mildly deleterious mutations (Muller 1950; Crow 1993; Manna et al. 2011).

These theoretical studies therefore suggest that deleterious mutations in themselves are unlikely to provide a mechanism for creating an initial selective advantage to autosomal inversions in a randomly mating population. Indeed, if a new inversion arises on a unique haplotype, the process of accumulation of mutational load within the inversion subpopulation will take a substantial amount of evolutionary time, and cannot contribute to any initial selective effect of the inversion. It is, however, an open question as to whether the fitness differences among karyotypes mentioned above could have a significant component resulting from the process proposed by Sturtevant and Mather (1938), whereby genetic drift causes the inverted and standard arrangements to differ in their genetic content. There is a strong, but not perfect, analogy with population subdivision, whereby genetic drift can cause local populations to diverge at weakly selected loci subject to mutation to deleterious variants. Provided that these mutations are at least partially recessive with respect to their fitness effects, interpopulation crosses may show heterosis, due to different loci having accumulated different deleterious mutations in different populations (Whitlock et al. 2000; Glémin et al. 2003; Roze and Rousset 2004; Spigler et al. 2017; Charlesworth 2018); such heterosis has indeed been observed in populations of animals and plants (reviewed in Charlesworth 2018). In addition, theory predicts that populations with a smaller effective size (*N*_*e*_) should allow the accumulation of deleterious mutations and hence acquire larger mutational loads than populations with a larger *N*_*e*_ (Wright 1931; Kimura et al. 1963; Bataillon and Kirkpatrick 2000; Charlesworth 2018).

Such differences in *N*_*e*_ can arise either from differences in the adult population size itself, differences in the mating system (especially the frequency of inbreeding), differences in recombination rates associated with different levels of genetic hitchhiking effects, or a combination of all three factors. Again, there is empirical support for this predicted effect of *N*_*e*_, both at the level of measurements of fitness components (Leimu et al. 2006; Lohr and Haag 2015; Charlesworth 2018) and population genomic indicators of the efficacy of selection against deleterious mutations, e.g., Robinson et al. (2022) (population size), Glémin et al. (2006) (mating system), and Campos et al. (2014) (recombination rate).

The purpose of the present paper is to investigate the properties of a population genetic model of mutation, selection and drift acting on a polymorphism for an autosomal inversion and a standard arrangement, in which the inversion polymorphism is maintained by sufficiently strong selection that the frequency of the inversion is constant over time and space. Similar assumptions were used in an earlier paper that examined purely neutral differentiation between the two arrangements (Charlesworth 2023). In order to simplify the calculations, and to maximise the effects of drift and mutation within arrangements, no recombinational exchange between arrangements in heterokaryotypes is allowed. The results should therefore provide upper bounds on the likely size of effects.

Models are developed of both a single, randomly mating population and of a population divided into a large number of local populations; it might be expected that population subdivision, with its greater opportunities for drift, would enhance divergence among arrangements at sites under selection. The model assumes a large number of freely recombining sites subject to selection and mutation, with a wide distribution of selection coefficients over sites; no attempt is made to investigate the consequences of enhanced Hill-Robertson interference among sites due to restricted recombination in heterokaryotypes, which was included in the simulations of the fates of autosomal inversions in Berdan et al. (2021) and Jay et al. (2022). Unless an autosomal inversion is very rare or very common, there should be sufficient recombination within homokaryotypes to prevent major Hill-Robertson effects; very low recombination rates are sufficient to prevent the operation of Muller’s ratchet (Charlesworth et al. 1993). Hill-Robertson effects are, however, likely to be greatly enhanced in systems like newly evolving sex chromosomes, where at least part of a proto-Y or proto-W chromosome is held permanently heterozygous over its counterpart and fails to recombine (Charlesworth and Charlesworth 2000), so these cases are not considered here.

The overall conclusion is that a low frequency arrangement will have a higher mutational load and exhibit weaker population genomic signals of purifying selection than its counterpart. Heterokaryotypic superiority in fitness is, however, unlikely to be observed unless the inverted and standard sequences are approximately equal in frequency, and it is likely to be small in magnitude unless the inversion contains millions of sites under selection. Population subdivision has only a small effect on the load and population genomic statistics.

## A model of the mutational load associated with an inversion polymorphism

### General considerations

The main symbols used in this paper are defined in Table 1. Consider first a single randomly mating, discrete generation population of size *N*, assuming a Wright-Fisher model of reproduction such that *N* is equal to the effective population size, *N*_*e*_. The frequencies of the two karyotypes, the inverted (*In*) and standard *(St*) arrangements are denoted by *x* and *y* = 1 − *x*; designation as *In* versus *St* is purely arbitrary in the situation considered here, so the convention that *x* ≤ ½ is used. Balancing selection on the inversion is assumed to be sufficiently strong that *x* can be treated as constant over time. Let *q*_*i*_ and *p*_*i*_ = 1 − *q*_*i*_, be the respective frequencies of the mutant (A_2_) and wild-type allele (A_1_) at a given nucleotide site within haplotypes carrying a type *i* karyotype, where *i* =1 corresponds to *In* and *i* = 2 to *St*. In a given generation, there will be random drift as well as selection within the populations of *In* and *St* karyotypes, so that in general *q*_1_ ≠ *q*_2_. Drift occurs independently within karyotypes, so that the effective population sizes of carriers of *In* and *St* are *N*_1_ = *Nx* and *N*_2_ = *Ny*, respectively.

**Table 1.** Definitions of the most important symbols used in the text.

|  |  |
| --- | --- |
| $x$ and $y$ | frequencies of the inversion ($In$) and standard arrangement ($St$), respectively |
| $N$ | size of the population (single population case) or size of a local population (subdivided population case) |
| $d$ | number of demes in the subdivided population case |
| $N_T = dN$ | sum of local population sizes for a subdivided population |
| $L_i$ | mutational load for the homokaryotype subpopulation of type $i$ |
| $L_{12}$ | mutational load for $In/St$ heterokaryotypes |
| $B_i$ | inbreeding load for subpopulation $i$ |
| $u$ and $v$ | mutation rates from wild-type to mutant alleles and vice-versa |
| $\alpha$ and $\beta$ | $u$ and $v$ scaled by $4N$ or $4N_T$, depending on context |
| $m$ and $M$ | migration rate and migration rate scaled by $4N$ |
| $\langle q_i \rangle$ | expected frequency of mutant alleles for subpopulation $i$ |
| $V_{q_i}$ | variance in frequency of mutant alleles for subpopulation $i$ |
| $s$ | selection coefficient against homozygotes for a deleterious mutation |
| $h$ | the dominance coefficient for heterozygotes for a deleterious mutation |
| $a$ | shape parameter of the gamma distribution of selection coefficients |
| $\bar{\gamma}$ | mean selection coefficient against a deleterious mutation, scaled by $2N$ or $2N_T$ |

Subscripts $i = 1$ and $i = 2$ are used to denote parameters applicable to the $In$ and $St$ subpopulations, respectively. Subscript $d$ is used to denote within-deme parameters.

The expectation of *q*_*i*_ is denoted by ⟨*q*_*i*_⟩, with ⟨*p*_*i*_⟩ = 1 − ⟨*q*_*i*_⟩, where the angle brackets denote an expectation taken over the probability distribution of *q*_*i*_. The variance in *q*_*i*_ in a given generation over the probability distribution generated by drift is denoted by *V*_*qi*_, with the corresponding *F* statistic (Wright 1951) given by *F*_*i*_ = *V*_*qi*_ /⟨*p*_*i*_⟩⟨*q*_*i*_⟩. In the absence of recombination but the presence of selection there will be a negative covariance *C*_12_ between *q*_1_ and *q*_2_, with a correspondingly negative correlation coefficient *R*_12_, since a higher frequency of the mutant allele in one karyotype results in a higher frequency of mutant homozygotes in *In*/*St* individuals, enhancing the strength of selection against the mutation in the other karyotype. For neutral sites in the absence of recombination, however, *C*_12_ = *R*_12_ = 0.

Following Kimura et al. (1963), equations can be written for the genetic loads at a single diallelic autosomal locus, assuming a homozygous selection coefficient *s* and dominance coefficient *h*. The fitness of mutant homozygotes relative to wild-type is 1 − *s* and the fitness of heterozygotes is 1 − *hs*; *s* may vary across loci, but *h* is treated as a constant, although it is easy to relax this assumption. *L*_*i*_ is the genetic load for individuals homozygous for karyotype *i* produced by random mating within the population, defined as the expected reduction below 1 of their mean fitness relative to wild-type homozygotes. *L*_12_ is the corresponding load for heterokaryotypes. The homozygous load *H*_*i*_ is the reduction below 1 in the expected relative fitness of individuals made homozygous for gametes with karyotype *i*, with probability ⟨*q*_*i*_⟩ of being homozygous for the mutant allele. *B*_*i*_ is the inbreeding load for karyotype *i*, defined as *H*_*i*_ *-L*_*i*_ (Charlesworth and Charlesworth 2010, p.173).

Following Charlesworth (2018), some simple algebra yields the following expressions for these quantities:

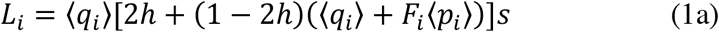

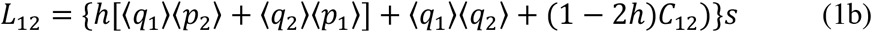

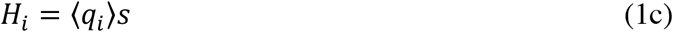

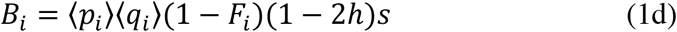

where:

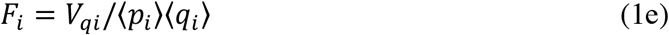

The selective difference between a heterokaryotype and a homokaryotype of class *i*, can be measured by *t*_*i*_ = *L*_*i*_ − *L*_12_. Heterozygote advantage exists if both *t*_*i*_ are positive. To obtain a convenient expression for this quantity, it is useful to rearrange Equations (1) by writing ⟨*q*_1_⟩ = ⟨*q*⟩ + ⟨*δq*⟩, ⟨*q*_2_⟩ = ⟨*q*⟩ − ⟨*δq*⟩, where 2⟨*δq*⟩ is the expected difference in the frequency of A_2_ between *In* and *St*. ⟨*δq*⟩ will be non-negative if *x* < ½, due to the greater effectiveness of drift relative to selection in a smaller population (Kimura et al. 1963). We then obtain the following expressions:

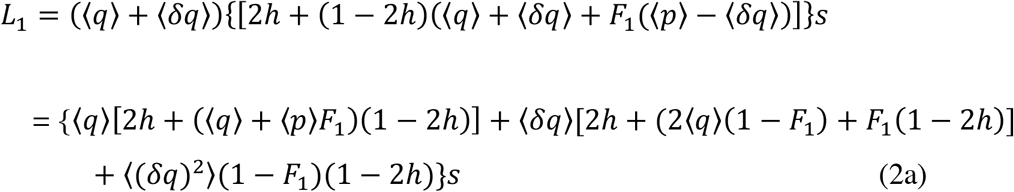

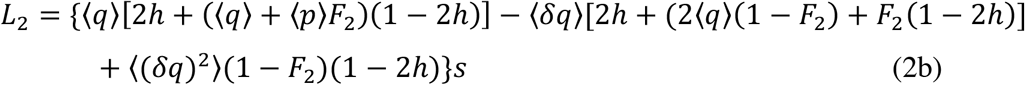

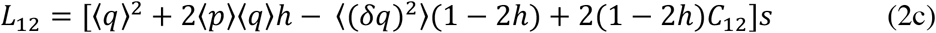

Equation (2c) shows that, when *h* < ½ the mean fitness of heterokaryotypes is increased by a difference in the expected frequencies of deleterious mutations between the two karyotypes, as expected intuitively.

These expressions yield the following results for the selective differences between heterokaryotypes and homokaryotypes:

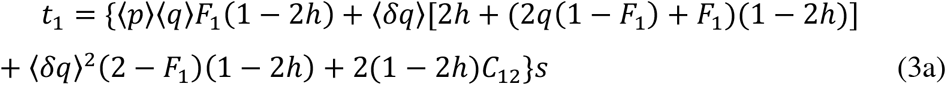

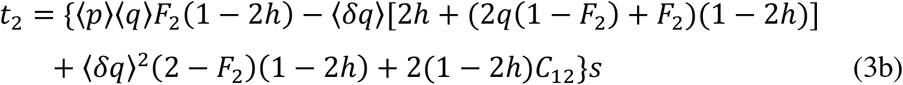

It is easily seen that, for an inversion with *x* < ½ and ⟨*δq*⟩ > 0 (see above), we have *t*_1_ > 0 provided that *h* ≤ ½ and *C*_12_ ≥ 0, so that *In*/*In* then has a lower fitness than *In*/*St*. The expectation for a low frequency inversion is thus that *t*_1_ > 0, although the magnitude of the effect is likely to be small for a single, large population. Population subdivision, causing both *F*_*i*_ to be > 0, will enhance *t*_1_. With *F*_1_ > 0 and *h* < ½, this can also be the case when ⟨*δq*⟩ = 0, reflecting the fact that population subdivision causes a reduction in mean fitness by increasing the frequencies of homozygotes; this effect is not experienced by *In*/*St* individuals unless *C*_12_ > 0, which can be ruled out by the argument given above.

If ⟨*δq*⟩ > 0 and *h* ≤ ½, the condition for *t*_2_ > 0 is more stringent than for *t*_1_ > 0, due to the opposite sign of the term in ⟨*δq*⟩ in Equations (3a) and (3b), especially if *F*_2_ is close to zero. In species with low levels of population subdivision, it may therefore be difficult to find conditions in which there is an advantage to *In*/*St* over both homokaryotypes, unless the inversion frequency is close to ½, so that ⟨*δq*⟩ ≈ 0.

These results can be generalised to the case of a subdivided population with a constant frequency of the inversion across all local populations (demes), by taking expectations of within-deme allele frequencies across populations, as described in section 2 of the Appendix.

### A single population: modeling drift and selection

In order to obtain numerical results for the load statistics described above, expressions for the means and variances of the *q*_*i*_, as well their covariance, are needed. Recombination is assumed to be absent in heterokaryotypes. We first consider the expected changes in allele frequencies due to selection within each karyotype. For *In* karyotypes, the marginal fitness of haplotypes carrying the wild-type allele (A_1_) at a locus is easily seen to be:

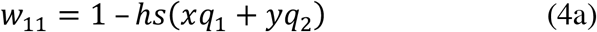

Similarly, the marginal fitness of *In* haplotypes carrying the mutant allele is:

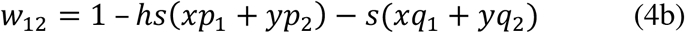

The net expected change in the frequency of A_2_ within *In* karyotypes due to selection (neglecting second-order terms in *s*) is thus:

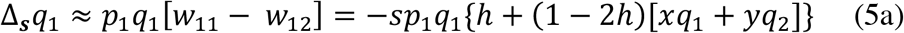

Similarly, the net expected change in the frequency of A_2_ within *St* karyotypes due to selection is:

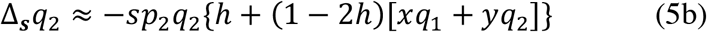

Equations (4) and (5) bring out the interdependence between the evolutionary processes in the two karyotypes when *h ≠* ½. To proceed further, the effects of mutation and drift also need to be analysed, such that an expression for the stationary joint probability density function (p.d.f.) for *q*_1_ and *q*_2_, *ϕ* (*q*_1_, *q*_2_), can be obtained. Kimura (1964, p.41) derived a pair of coupled forward diffusion equations describing the joint stationary p.d.f. for two variables, using the principle that a zero flux of the probability density of each variable across all their values guarantees a stationary joint distribution. This method can be applied to the above selection equations, together with the terms arising from mutation.

Let the rates of mutation from A_1_ to A_2_ and vice-versa be *u* and *v*, respectively, with scaled mutation rates *α*_1_ = 4*N*_1_*u, α*_2_ = 4*N*_2_*u, β*_1_ = 4*N*_1_*v, β*_2_ = 4*N*_2_*v*. The mutational bias towards deleterious mutations, *k*, is equal to *u*/*v*. As shown by Kimura (1964), in order to analyse this type of situation it is most convenient to use the natural logarithm of *ϕ, ψ* = ln *ϕ*. If drift affects *q*_1_ and *q*_2_ independently, as is the case here, the conditions for a stationary distribution are:

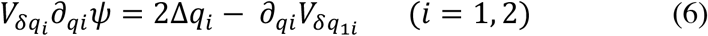

where 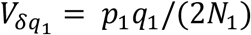 and 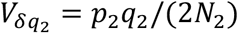 are the variances in the changes in allele frequencies per generation due to drift within *In* and *St*, respectively. Correspondingly, 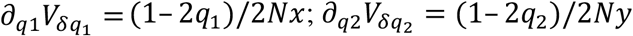. The Δ*q*_*i*_ are given by the selection equations (4) and (5) together with the relevant mutation terms.

For a meaningful solution of Equation (6) to exist, *ψ* must have an exact differential, 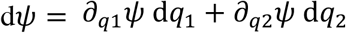, which requires 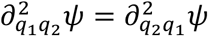 (Kimura 1964, p.41). Kimura showed that this is the case for the mutation terms, so we need only consider the selection terms contributed by Equations (4) and (5). The following expression satisfies both this condition and Equation (6) for the selection contribution to *ψ*:

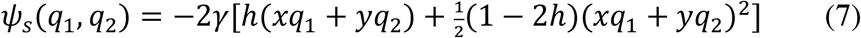

where *γ* is the scaled selection coefficient, 2*Ns*.

The full solution to Equation (6) is thus:

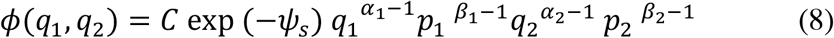

where *C* is a normalisation constant, given by the inverse of the double integral of *φ* over the closed intervals (0, 1) with respect to *q*_1_ and *q*_2_.

### A single population: obtaining the mutational load and population genomic statistics

To obtain the single locus load statistics for a given *h*, numerical integration of Equation (8) and its product with powers and crossproducts of the *q*_*i*_ can be performed, for the purpose of determining the expectations and variances of the *q*_*i*_ and their covariance *C*_12_. The corresponding *F*_*i*_ statistics can be obtained from Equation (1d). The means of the single locus load statistics, with the terms in *s* omitted, can then be obtained using Equations (1). Details of the integration procedures are given in section 1 of Supplementary File S1.

In order to calculate the load statistics themselves under reasonably realistic assumptions, a gamma distribution of the scaled selection coefficient *γ* = 2*Ns* is assumed, with a p.d.f. given by:

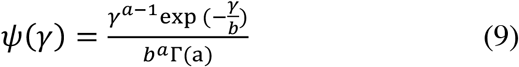

where *a* is the shape parameter, 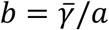 is the location parameter and Γ (*a*) is the gamma function. This distribution has been widely used in population genomic methods for estimating of the distribution of fitness effects of deleterious mutations (e.g., Booker et al. 2017).

The values of 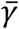 and the shape parameter *a* are chosen to correspond to estimates from population genomics studies just mentioned, which indicate 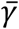 values of hundreds or thousands for nonsynonymous mutations and shape parameters of approximately 0.3, implying a wide distribution of selection coefficients. It is assumed that fitness effects are multiplicative across sites, so that the products of the expectations of Equations (1a) − (1c) over the distribution of *γ* with the number of sites, *n*_*s*_, correspond to the natural logarithms of the corresponding multi-site load statistics. The exponentials of the negatives of these expressions then yield the mean fitnesses of the karyotypes concerned, relative to that of mutant-free individuals. The exponential of the negative of the product of *n*_*s*_ and the expectation of Equation (1c) yields the fitness of totally inbred individuals of karyotype *i* relative to outbred individuals that are homozygous for karyotype *i*, i.e., a measure of the inbreeding depression experienced by carriers of karyotype *i*.

The net selection coefficient against homozygosity for karyotype *i* relative to heterokaryotypes for a given *n*_*s*_ is measured by:

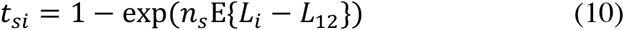

where E{} indicates the expectation over the distribution of *γ* (as opposed to an expectation over the distribution of *q*, denoted by angle brackets)

It is also of interest to summarise the expected patterns of variation at the individual loci. To do this, the p.d.f. for *q*_1_ and *q*_2_ can be used to calculate the expected nucleotide site diversities within each karyotype subpopulation as *π*_*i*_ = 2⟨*p*_*i*_⟩⟨*q*_*i*_⟩(1 − *F*_*i*_) for a given selection coefficient, which are then averaged over the distribution of *s*. In addition, the expected proportion of segregating sites for a sample of *n* alleles, *S*_*ni*_, can be determined as described in section 2 of Supplementary File S1. Division by the sum of the harmonic series, 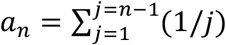, yields the expected values of Watterson’s theta (*θ*_*wni*_) for each subpopulation (Watterson 1975). The skew of the distribution of segregating variants towards rare variants for subpopulation *i* can conveniently be measured by Δ*θ*_*wni*_ = 1 − *π*_*i*_/*θ*_*wi*_ (Campos and Charlesworth 2019).

For practical purposes of computation, it is convenient to divide the range of values of *γ* into several zones according to the strength of selection and to compute the integrals of the load statistics over each zone separately. The overall values of the load statistics are then given by summing the results over all zones. The details are given in section 1 of the Appendix, and the computer code for generating the numerical results for this case is given in Supplementary File S2.

### A finite island model metapopulation

In this case, a metapopulation of total size *N*_*T*_ is divided into a large number *d* of subpopulations (demes), each of size *N* = *N*_*T*_ /*d*. A Wright-Fisher model is assumed to apply to each deme. *N* is assumed to be sufficiently large that the frequency of the inversion is held at the same frequency *x* in all demes. A fraction *m* of each deme is derived by migration from a pool with equal contributions from all demes. Let the current mean frequency across all demes of the mutant allele A_2_ at a locus be 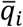 for karyotype *i*, so that migrants contribute 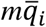 to the new frequency of A_2_ among type *i* haplotypes within a deme and 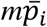 to the new frequency of A_1_.

This model poses the problem that the evolutionary processes within demes change the values of the 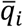, so that they cannot realistically be treated as fixed quantities. Following previous treatments of this problem, it is assumed here that the process of change in the 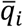 can be described by a pair of coupled diffusion equations, using the expectations of *p*_*i*_*q*_*i*_/ (2*N*_*T*_) over all *d* demes as the drift variance terms together with the corresponding expectations of the expressions for the deterministic changes in 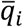 (Whitlock 2002; Cherry and Wakeley 2003; Roze and Rousset 2003;

Wakeley 2003). The mutational contributions to the latter are simply obtained by substituting 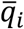 for *q*_*i*_ in the standard formulae for the deterministic changes in the 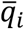, since the mutational changes are linear in *q*_*i*_. The expectation of *p*_*i*_*q*_*i*_ over demes conditioned on 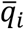 can be written as 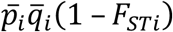, where *F*_*STi*_ is the variance among demes in *q*_*i*_ divided by 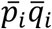. The total effective population size for the metapopulation for karyotype *i, N*_*emi*_, is thus equal to *N*_*T*_/(1 − *F*_*STi*_) (Wakeley and Aliacar 2001).

The non-linearity with respect to *q*_*i*_ of the selection terms for allele frequency changes within demes (after division by *p*_*i*_*q*_*i*_) when *h* ≠ 0.5 means that an exact closed expression for their contributions to the expected changes in the 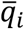 cannot be obtained, except in the absence of dominance (Whitlock 2002; Roze and Rousset 2003; Wakeley 2003). However, a useful approximation can be obtained by neglecting the third moments about their means of the within-deme allele frequencies; these moments are necessarily smaller than the variances of the 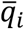, and will be considerably smaller when selection is sufficiently strong in relation to drift that the 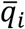 are close to zero. When selection is sufficiently weak, a neutral approximation for the third moment can be used (Whitlock 2002), but will in general over-correct when selection is strong and so is not employed here. The error introduced by ignoring this correction affects only a small portion of the distribution of selection coefficients where *s* is *O*(1/2*N*_*T*_) and is thus unlikely to be important for the load and population genomic statistics calculated here.

The following expressions are obtained after some algebra:

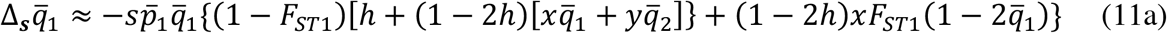

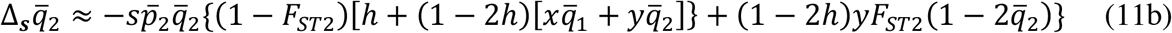

We also have the following expression for the variance in 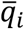 due to drift, e.g., (Whitlock 2002):

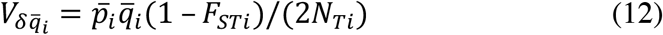

where *N*_*T*1_ = *N*_*T*_*x* and *N*_*T*2_ = *N*_*T*_*y*.

Combining Equations (11) and (12), using the approach that led to Equations (8) and carrying out some rearrangements of terms, we obtain the equivalent of the *ψ* function for the panmictic case, which describes the selection contribution to the logarithm of the p.d.f. for the metapopulation, writing *γ*_*m*_ = 2*N*_*T*_*s* for the scaled selection coefficient with regard to the whole metapopulation and *G*_*i*_ = *F*_*STi*_/(1-*F*_*STi*_):

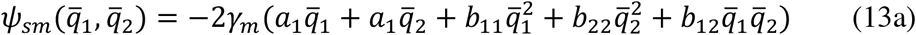

where

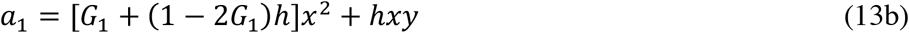

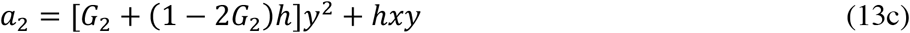

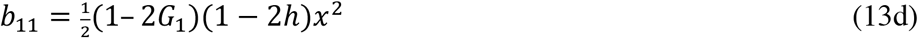

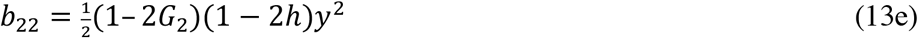

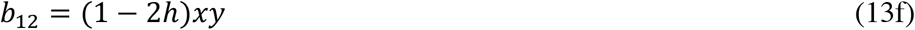

When selection is sufficiently weak in relation to migration that 2*N*_*T*_*s* << *M*, where *M* is the scaled migration parameter 4*Nm*, we have *F*_*ST1*_ ≈ 1/(1 + *Mx*) and *F*_*ST2*_ ≈ 1/(1 + *My*),. For sites independent of the inversion, we have *F*_*STn*_ ≈ 1/(1 + *M*). Purifying selection is expected to reduce the *F*_*STi*_ below their neutral values (Charlesworth and Charlesworth 2010, p.353), so that use of the neutral approximations overestimates the effect of drift relative to selection. An approximate correction for the effect of selection on *F*_*STi*_ is used here, based on Equation (B7.8.4) of (Charlesworth and Charlesworth (2010, p.354); the term 4*Nhs* is added to *M* in the formula for *F*_*ST*_. This procedure is only rigorous for *Nhs* >> 1; numerical studies show that it very slightly reduces the effect of population subdivision compared with using the purely neutral expression (Supplementary Table 1).

Combining Equations (13) with the mutational terms, the bivariate p.d.f. has a similar form to Equation (8), except that 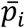 and 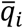 are used instead of *p*_*i*_ and *q*_*i*_, and the scaled mutational parameters involve the products of 4*N*_*T*1_*/*(1 − *F*_*ST*1_) and 4*N*_*T*2_*/*(1 − *F*_*ST*2_) with the relevant mutation rates. Thus, *α*_*i*_ and *β*_*i*_ are replaced by *α*_*im*_ = 4*N*_*Ti*_ *u/*(1 − *F*_*STi*_) and *β*_*im*_ = 4*N*_*Ti*_ *v/*(1 − *F*_*STi*_), respectively.

In order to apply Equations (1), it is necessary to consider the p.d.f. of allele frequencies within demes. Following Cherry and Wakeley (2003) and Wakeley (2003), conditioning on the current values of the 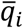 for a deme with frequencies *q*_*i*_ we have the general expression:

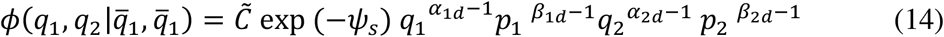

where *ψ*_*s*_ has the same form as in Equation (7), 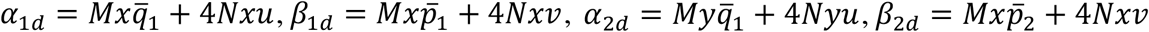 (the subscript *d* is used to indicate within-deme values).

An exact calculation of the load statistics for a given *s* would require multiplication of this function by the bivariate p.d.f. for the 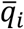, followed by integrations over the *q*_*i*_ and 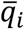 to obtain the requisite quantities to insert into Equations (1); this would then be followed by integration over the distribution of *s*. In order to avoid this cumbersome procedure, partitioning of the distribution of *s* into different zones was used together with some approximations, similarly to what was done for the case of a single panmictic population. The details are given in section 2 of the Appendix. The computer code for generating the numerical results for this case is given in Supplementary File S3.

## Results

### General considerations

Intuitively, the mutational load associated with each homokaryotype in an inversion polymorphism would be expected to be strongly affected by the dominance coefficient (*h*), the scaled strength of selection (2*Ns* = *γ*), the mutational rate to deleterious mutations (*u*), and the frequency of the inversion (*x*). Dominance coefficients less than one-half are well known to be required for inbreeding depression and heterosis, whose magnitudes are inversely related to *h* (Charlesworth and Charlesworth 2010, Chap. 4). It would thus be expected than the reduction in fitness associated with an arrangement, and any fitness advantage to *In*/*St* heterokaryotypes, would decrease with *h* but increase with *u*, if *γ* is sufficiently small that deleterious mutations are significantly affected by drift.

It is less clear how these properties are related to *γ*, since stronger selection reduces the frequencies of deleterious mutations but also reduces their effects on fitness if they rise to high frequencies. However, if selection is so strong in relation to drift that allele frequencies are at mutation-selection equilibrium, no differentiation in allele frequencies between *In* and *St* will occur, removing any possibility of a selective advantage to heterokaryotypes (Sturtevant and Mather 1938; Nei et al. 1967), and the mean fitnesses of both homokaryotypes with *n*_*s*_ selected sites will be equal to the deterministic value, exp(−2*n*_*s*_*u*) ≈ 1 -2*n*_*s*_*u* when fitnesses are multiplicative and *h* is > 0 (Haldane 1937).

The rarer of the two arrangements experiences more genetic drift than its counterpart, so that *x* < 0.5 means that inversion homokaryotypes should have a lower overall mean fitness than standard homokaryotypes. It is less clear when an advantage to heterokaryotypes can be generated; the increased load associated with the rarer arrangement may simply generate a net selective advantage to its counterpart, especially when mutations are only partially recessive. Finally, it is likely that population subdivision will increase the magnitude of the mutational loads for each homokaryotype and the selective differences among the three karyotypes, because drift within demes in a metapopulation occurs at a faster rate than in a single population with the same size as the metapopulation. However, this is counterbalanced by a slower rate of drift for the population as a whole (Whitlock 2002; Cherry and Wakeley 2003; Wakeley 2003), so the net effect is hard to predict intuitively.

### A single population: load statistics

Numerical results for a single randomly mating population are presented here. These are based on Equations (1) for individual selected sites together with the procedures for combining the effects of mutation, drift and selection for all sites that were described above. The number of sites (*n*_*s*_) was set to 10^5^, corresponding to an inversion containing 100 genes with a mean of 1000 nonsynonymous sites per gene. The expectations of the single-locus mutational loads (*L*_*i*_ and *L*_12_) and the inbreeding loads (*B*_*i*_), as defined by Equations (1), are then multiplied by 10^5^ to obtain their net values. If multiplicative fitnesses are assumed, these quantities are equivalent to the negatives of the natural logarithms of the corresponding mean fitnesses (for the *L’*s) or differences in log mean fitnesses (for the *B’*s). The values of the corresponding mean fitnesses for a different number of sites, *n*_*s*1_, can be found by taking the exponentials of their negatives multiplied by *n*_*s*1_/10^5^. The selection coefficients *t*_*i*_ against the two homokaryotypes are obtained by exponentiation of the product of *n*_*s*1_ and the expectation of (*L*_*i*_ − *L*_1*2*_) (Equation 10). If these are small, as is the case in practice, *t*_*i*_ ≈ *n*_*s*1_ E{*L*_*i*_ − *L*_12_}. Heterokaryotype advantage requires both of the *t*_*i*_ to be positive; if one is positive and the other negative, there is directional selection in favour of the karyotype with the negative *t*_*i*_.

Figure 1 displays the values of the relevant load statistics and the selection coefficients against homokaryotypes as a function of the dominance coefficient *h*, for three different inversion frequencies and two different mean scaled selection coefficients (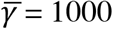 and 4000), using a gamma distribution of selection coefficients with shape parameter *a* = 0.3, and a mutation rate to deleterious alleles of *u* = 5 x 10^−9^. These values are broadly consistent with population genomic estimates from *Drosophila melanogaster* (Charlesworth 2015). A strong mutational bias to deleterious mutations of *k* = 1.5 was assumed, in order to maximise the magnitude of the mutational loads and selection coefficients.

**Figure 1.**
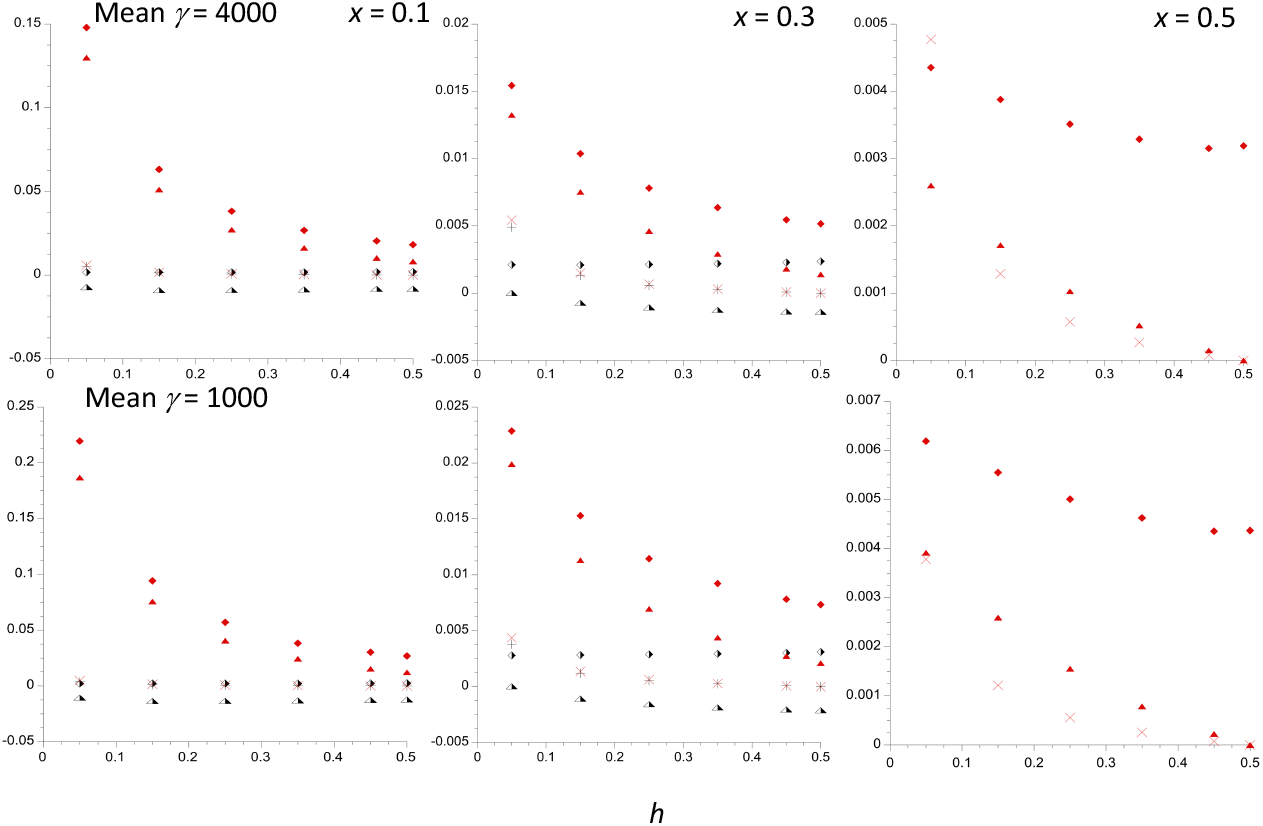
The mutational load statistics for a single population of size *N* = 10^6^ with 10^5^ selected sites, plotted against the dominance coefficient *h*. The results for three different frequencies of the inversion are shown. The selection coefficients follow a gamma distribution with a mean of 5 x 10^−4^ and a shape parameter of 0.3. The mutation rate to deleterious alleles is 5 x 10^−9^ per basepair, with a mutational bias towards deleterious variants of 1.5. The upper and lower panels have product of 2*N* and mean *s* of 4000 and 1000, respectively. The filled and half-filled symbols denote values for the *In* and *St* subpopulations, respectively. The lozenges are the net mutational loads within the respective karyotypes, and the crosses are the corresponding inbreeding loads. The triangles are the selection coefficients against homokaryotypes relative to the heterokaryotype. Only the values for *In* are shown when *x* = 0.5.

If there were no effects of drift, the net load (*L*) for both subpopulations would be approximately 2*n*_*s*_*u* = 0.001, and the net inbreeding load (*B*) would equal *n*_*s*_*u*(*h*^-1^ − 2); this gives *B* = 0.002 with *h* = 0.25 and 0.036 with *h* = 0.05) − see Charlesworth and Charlesworth (2010, Chap. 4). The selection coefficients on both karyotypes would be zero. The numerical results shows that the loads for both *In* and *St* subpopulations are always substantially larger than the deterministic values, even for semi-dominant mutations (Supplementary Table 2). With *x* = 0.1 and 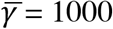, the load for *In* (*L*_1_) is more than twenty-fold greater than the deterministic value for all *h* values between 0.05 and 0.5. Both *L* values are higher for the lower 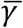 values, as might be expected from the larger effects of drift in this case. Similarly, with *x* < ½, *L*_1_ is always larger than *L*_2_, and decreases with *x* for a given *h* and 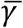, even for *h* = ½; in addition, *L*_1_ strongly decreases with *h* for *x* = 0.1 and 0.3, whereas *L*_2_ is only slightly affected by *h*. The fact that *L*_1_ < *L*_2_ with *h* = ½ and *x* < ½ shows that the increase in load due to smaller population size is partly caused by increased mean frequencies of deleterious mutations, not simply by increased frequencies of homozygotes (see Figure 4).

In contrast, the *B*_*i*_ values are always lower than the deterministic values, reflecting the reduction in variability caused by drift, and are decreasing functions of *h* (vanishing when *h* = 0.5), although this is hard to see from Figure 1 due to their overall very small values. These effects are especially noticeable for *B*_1_ when *x* = 0.1 and 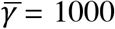 = 1000. The small *B*_*i*_ values reflects the fact that, except for the lowest dominance coefficient (*h* = 0.05), the *L*_*i*_ are always quite close to the corresponding homozygous loads (the *H*_*i*_ of Equation 1c). For example, with 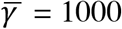 and *x* = 0.1, for *h* = 0.05, we have *L*_1_ = 0.212, *H*_1_ = 0.224, *L*_2_ = 0.002, *H*_2_ = 0.006; for *h* = 0.25, we have *L*_1_ = 0.057, *H*_1_ = 0.058, *L*_2_ = 0.002, *H*_2_ = 0.003.

There can be substantial selection against the inversion homokaryotypes at the lower *h* values, especially for the smaller *x* values: *t*_1_ reaches 0.19 with *h* = 0.05 and 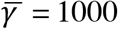, decreases sharply with *x* and *h*; it is only 0.04 for *h* = 0.25, *x* = 0.1 and 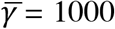. The values for 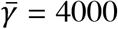 = 4000 are somewhat smaller, consistent with smaller effects of drift in causing divergence between the two subpopulations. In contrast, *t*_2_ is mostly negative and quite small (approximately -0.01 for a wide range of *h* values when *x* = 0.1 and 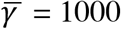), indicating weak directional selection against the inversion. Only when *x* is close to 0.5 does the heterokaryotype experience a slight advantage over both homokaryotypes (a maximal value of *t*_1_ = *t*_2_ ≈ 0.004 for *h* = 0.05 and 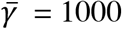 = 1000). With *x* = 0.5, the selective advantage to heterokaryotypes declines sharply with *h*, and is only 0.0016 for *h* = 0.25 and 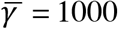; it vanishes when *h* = 0.5. With one million instead of 10^5^ selected sites, the heterokaryotype advantage would be approximately ten times as large. With *x* = 0.5, symmetry implies that *t*_1_ = *t*_2_, and the equality of effective population sizes means that the mean fitness of the heterokaryotypes is superior to that of the homokaryotypes purely because they have a lower frequency of mutant homozygotes than either of the homokaryotypic subpopulations.

It is also of interest to examine the effect of differences in population size and selection coefficients on the load statistics for a constant scaled selection strength. Figure 2 show results that are comparable to those in Figure 1, but with population sizes of 2 x 10^6^ and 10^5^, and a mean selection coefficient (0.001) that is one-half of that in Figure 1. A comparison of the top panels of the two figures shows that reducing the mean selection coefficient by one-half, but keeping 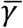 constant, results in a reduction in the *L* values, by a factor of close to 2 for the case of the *In* subpopulation with *x* = 0.1, but by considerably less for the *St* population, where *L*_2_ is only slightly greater than the deterministic value of 0.001, even for *h* = 0.05. The relative effect on a subpopulation is reduced as it increases in size, as might be expected intuitively. These effects reflect the fact that the load caused by drift is greater when the selection coefficients involved are larger, in contrast to the deterministic formula *L* = 2*u*. The two selection coefficients on homokaryotypes relative to heterokaryotypes, *t*_*i*_, show a similar pattern. In contrast, the inbreeding loads are only slightly reduced by a reduction in the strength of selection with constant 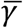. Further results for different deme sizes are shown in Supplementary Table 3.

**Figure 2.**
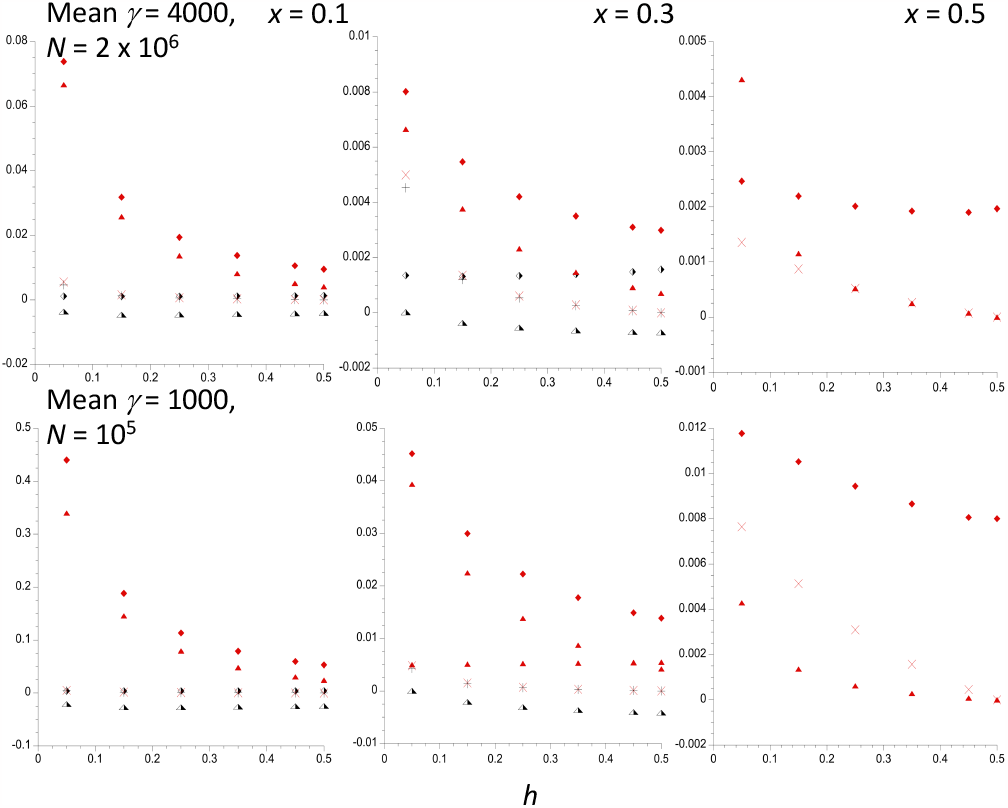
The mutational load statistics for a single populations of size *N* = 2 x 10^6^ (upper panel) and *N* = 10^5^ (lower panel), with the same mean selection coefficient (0.001), plotted against the dominance coefficient *h*. The results for three different frequencies of the inversion are shown. The other parameters are the same as in Figure 1. The filled and half-filled symbols denote values for the *In* and *St* subpopopulations, respectively. The lozenges are the net mutational loads within the respective karyotypes, and the crosses are the corresponding inbreeding loads. The triangles are the selection coefficients against homokaryotypes relative to the heterokaryotype. Only the values for *In* are shown when *x* = 0.5.

The effect of a reduced population size while holding mean *s* constant can be seen by comparing the upper and lower panels of Figure 2. As might be expected from the greater effects of drift in a smaller population, the *L*_*i*_ and *t*_*i*_ are considerable larger when the population size is reduced by a factor of twenty, especially for the inversion subpopulation with *x* = 0.1, for which *N*_1_ = 10,000 in the lower panel. However, with *x* = 0.5 and *h* = 0.25, the selective advantage to the heterokaryotypes is only 0.003 for the smaller population size. The inbreeding loads are barely affected by the population size difference.

### A single population: population genomics statistics

It is also of interest to examine the effects of an inversion polymorphism on population genomics statistics that can be used to assess the effects of the differences in effective population size between the *In* and *St* subpopulations, and between these and the part of the genome that is independent of the inversion polymorphism. Figure 3 shows the results for the *St* population using the same parameter values as in Figure 1 for three different values of *x*; the case with *x* = 0.1 is nearly equivalent to the situation for the rest of the genome. It can be seen, somewhat surprisingly, that the mean frequency of deleterious alleles in the *St* subpopulation (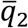, half-filled triangles), which includes both fixed and segregating sites, is nearly independent of *h*. It increases slightly as the size of the *St* population, *N*_2_ = *N*(1 − *x*), decreases as *x* changes from 0.1 to 0.5; the effect of *N*_2_ is somewhat greater for 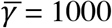, as would be expected from the greater effects of drift in this case. For example, with *h* = 0.25 and 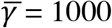, 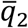 changes from 0.044 to 0.083 as *x* goes from 0.1 to 0.5, but only changes from 0.044 to 0.048 for 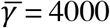 = 4000. This difference partly reflects the higher proportion of sites that can become fixed for deleterious mutations with weaker selection, as well as their higher mean frequency at segregating sites. Its behavior implies that the effect of population size on the load for a subpopulation expectation when the mean strength of selection is strong is largely caused by changes in the variance of *q*_2_ rather than its.

**Figure 3.**
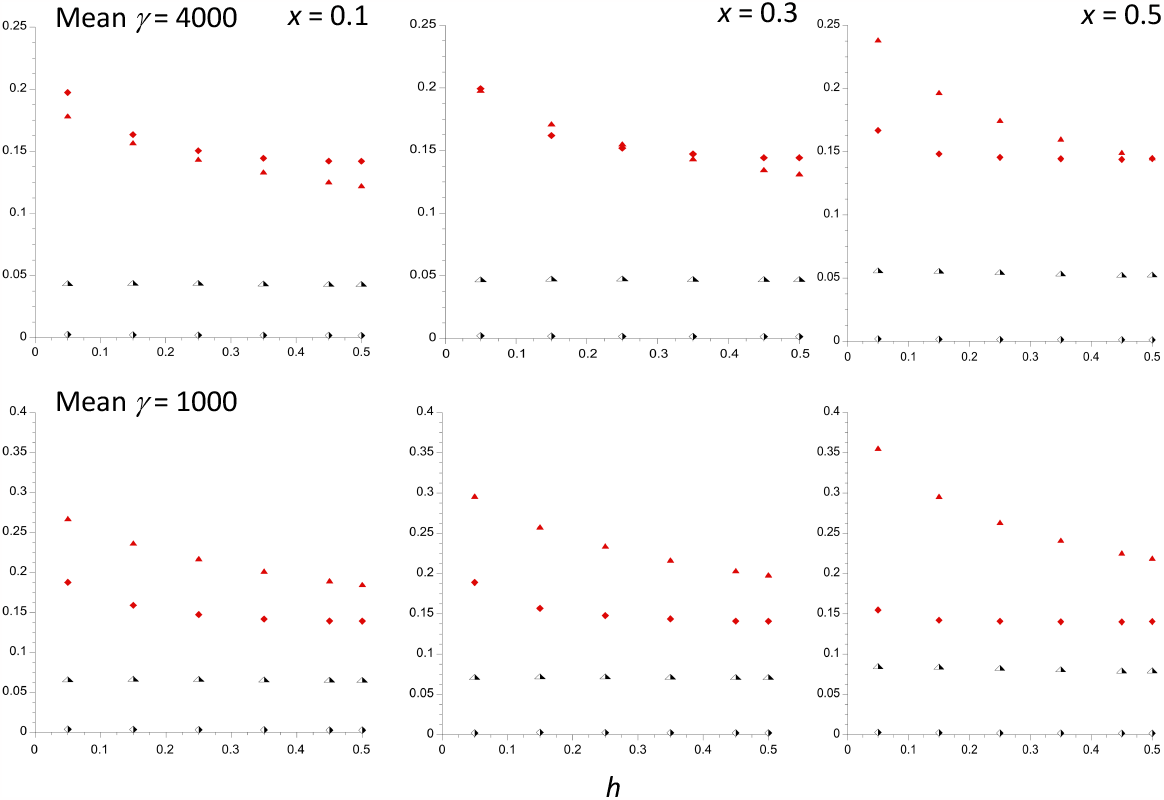
The population genomics statistics for the *St* subpopulation, plotted against the dominance coefficient *h*. The evolutionary parameters are the same as in Figure 1. The results for three different frequencies of the inversion are shown. The half-filled triangles are the mean frequencies of A_2_ and the half-filled lozenges are the mean diversities at selected sites. The filled triangles are the ratios of mean diversities at selected sites to mean diversities at neutral sites. The filled lozenges are the values of Δ*θ*_*w*_ for the selected sites.

**Figure 4.**
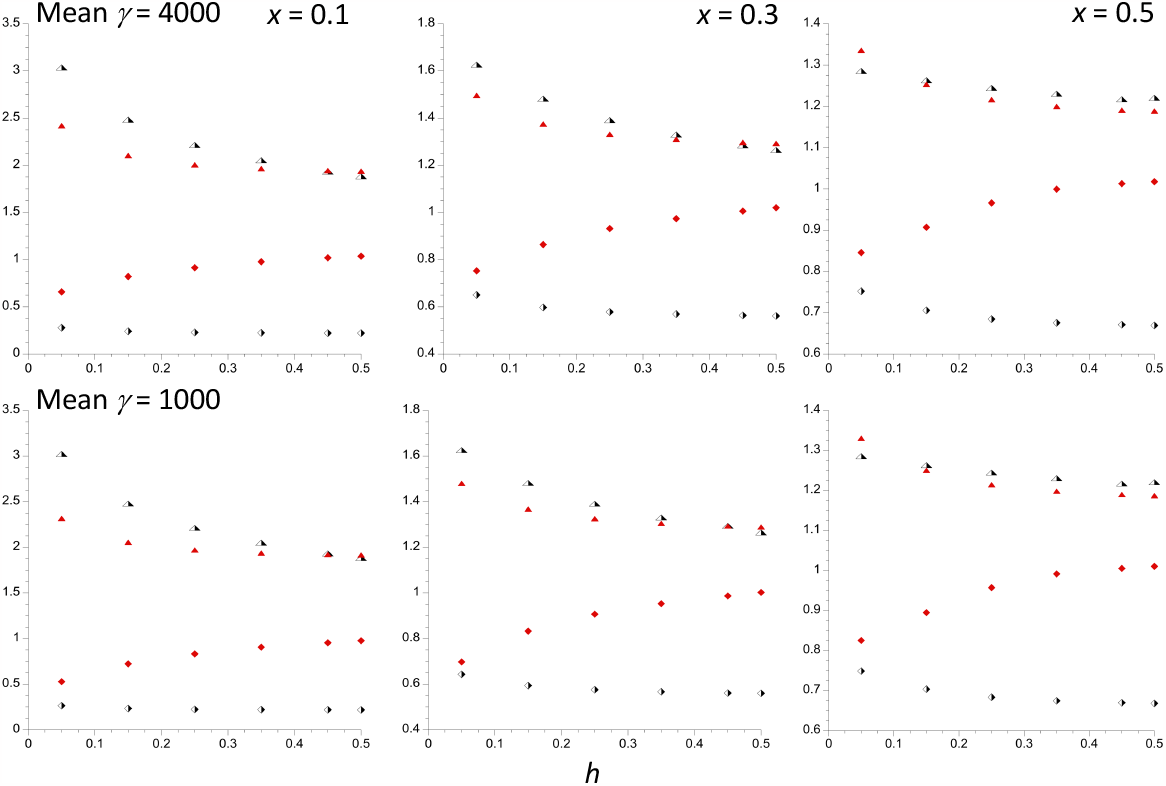
The first two panels in each row show the ratios of the population genomics statistics for the *In* subpopulation to those for the *St* subpopulation, plotted against the dominance coefficient *h*. The third panel shows the ratios of these statistics for the *St* subpopulation with an inversion frequency of 0.5 to their values for a frequency of 0.1, for which the corresponding ratio of population sizes is 0.556. The evolutionary parameters are the same as in Figure 1. The half-filled triangles are the ratios of the mean frequencies of A_2_; the half-filled lozenges are the ratios of the mean diversities at selected sites. The filled triangles are the ratios of mean diversities at selected sites to mean diversities at neutral sites. The filled lozenges are the ratios of Δθ_*w*_ values for the selected sites.

The mean nucleotide site diversity at the selected sites for *St* (*π*_*sel* 2_, half-filled lozenges) decreases somewhat with increasing *h* and decreasing *N*_2_, but is always between 0.0011 and 0.0025 for 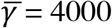 and between 0.0017 and 0.0037 for 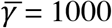. The ratio of *π*_*sel*2_ to the mean nucleotide site diversity at neutral sites (*π*_*sel*2_/*π*_*neut*2_, filled triangles) decreases with *N*_2_, especially when *h* is small. The measure of skew towards low frequency variants at selected sites, Δ*θ*_*w*2_, for a sample of 20 haploid genomes (filled lozenges) increases with *N*_2_, but the effect is weak unless *h* is small; the effect of *h* on Δ*θ*_*w*2_ is smaller than for *π*_*sel*2_/*π*_*neut*2_, and plateaus around *h* = 0.25 (for the definition of Δθ_*w*_, see the section *A single population: modeling drift and selection*).

These effects of *h* and *N*_2_ can also be seen in the right hand panels of Figure 4 (labelled as *x* = 0.5) which plot the ratios of 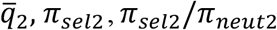 and Δ*θ*_*w*2_with *x* = 0.5 to their values with *x* = 0.1, a 1.8-fold difference in *N*_2_. The plots for *π*_*sel 2*_/*π*_*neut2*_ bring out clearly that a lower subpopulation size is associated with larger *π*_*sel 2*_/*π*_*neut2*_, especially when *h* is small. For Δ*θ*_*w*2_, the ratio is approximately 0.85 for *h* = 0.05, but increases rapidly towards 1 as *h* increases. The four-fold difference in 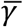 between the upper and lower panels has a remarkably small effect on the ratios for all four statistics.

The other two panels of Figure 4 display plots against *h* of the ratios of the same set of statistics for the *In* subpopulation to their values for the *St* subpopulation, for *x* = 0.1 and 0.3. The ratios of population sizes for *St* versus *In* are 9-fold for *x* = 0.1 and 2.3-fold for *x* = 0.3, flanking the ratio *N*_2_/ *N*_1_ for the right-hand panels. *N*_2_ for *x* = 0.3 is 1.4-fold greater than *N*_2_ for *x* = 0.5, which enhances the contrast between *In* and *St*. Accordingly, the patterns are more marked than for the right-hand-most panels, and are strongest for the case with *x* = 0.1, with a maximal ratio of *π*_*neut*1_/*π*_*neut*2_ to *π*_*sel 2*_/*π*_*neut2*_ of 2.4 at *h* = 0.05 and 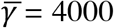. This is still much less than the ratio of 9 for neutral diversity. The ratio Δ*θ*_*w*1_ / Δ*θ*_*w*2_ with *x* = 0.1 is close to 0.5 for both 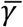 values when *h* = 0.05, but rapidly becomes close to 1 as *h* increases − for *x* = 0.1, *h* = 0.25 and 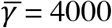, we have Δ*θ*_*w*1_ / Δ*θ*_*w*2_ = 0.915. The skew in the site frequency spectrum at selected sites is thus unlikely to be a powerful statistic for detecting a reduced efficacy of selection on a low frequency inversion.

### A subdivided population

As described in the section *General considerations, A finite island model metapopulation*, this case assumes that a metapopulation of total size *N*_*T*_ is divided into a large number *d* of demes, each of size *N* = *N*_*T*_ /*d*. A Wright-Fisher model applies to each deme, and the deme size is assumed to be sufficiently large that the frequency of the inversion is held at the same frequency *x* in all demes. A fraction *m* of each deme is derived by migration from a pool with equal contributions from all demes. For sites that are independent of the inversion, the level of neutral genetic differentiation between demes is described by *F*_*STn*_ ≈ 1/(1 + *M*), where *M* is the scaled mutation rate 4*Nm*.

Different levels of subdivision are characterized by different values of *F*_*STn*_; for simplicity, the subscript *n* is dropped in what follows. The scaled selection parameter *γ* is now defined as 2*N*_*T*_*s*, and the scaled mutation parameters are *α* = 4*N*_*T*_*u* and *β* = 4*N*_*T*_*v*.

Figure 5 shows the effects of population subdivision on the load statistics of most interest, together with the ratio of the diversities at selected sites for the *In* versus the *St* subpopulations, for the case of an inversion frequency of 0.1 in a metapopulation of 200 demes with a total size *N*_*T*_ = 10^6^ (*N* = 5000), and the same selection and mutation parameters as in Figure 1. The results for *F*_*ST*_ = 0 were obtained from the single population calculations described above. The results can be summarized very simply: there is a remarkably small effect of population subdivision as *F*_*ST*_ changes from 0 to 0.25, with the most marked effect occurring over the change from *F*_*ST*_ = 0 to *F*_*ST*_ = 0.05, especially for the smallest dominance coefficient (*h* = 0.05). For example, with 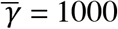 and *h* 0.05, the selection coefficients against the *In* and *St* homokaryotypes relative to the heterokaryotype change from *t*_1_ = 0.1868 and *t*_2_ -0.0107 with *F*_*ST*_ = 0 to 0.211 and − 0.0120 with *F*_*ST*_ = 0.05, reaching 0.250 and -0.0142 at *F*_*ST*_ = 0.25. With *h* = 0.25, the changes are much smaller: *t*_1_ = 0.0405 and *t*_2_ -0.0136 at *F*_*ST*_ = 0, and 0.0442 and -0.0146 at *F*_*ST*_ = 0.25. Both the absolute values of the loads and selection coefficients and their dependence on *F*_*ST*_ are much smaller with 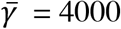 than 1000. As in the case of a single population, a selective advantage to the heterokaryotype is not found unless there are nearly equal frequencies of *In* and *St* (Supplementary Table 4). The magnitude of such an advantage is not greatly increased by subdivision; for example, with *x* = 0.5, 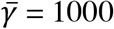 and *h* = 0.05, *t*_1_ = *t*_2_ = 0.0039 at *F*_*ST*_ = 0.0, and 0.00860 at *F*_*ST*_ = 0.25; with *h* = 0.25, the corresponding values are 0.00160 and 0.00196.

**Figure 5.**
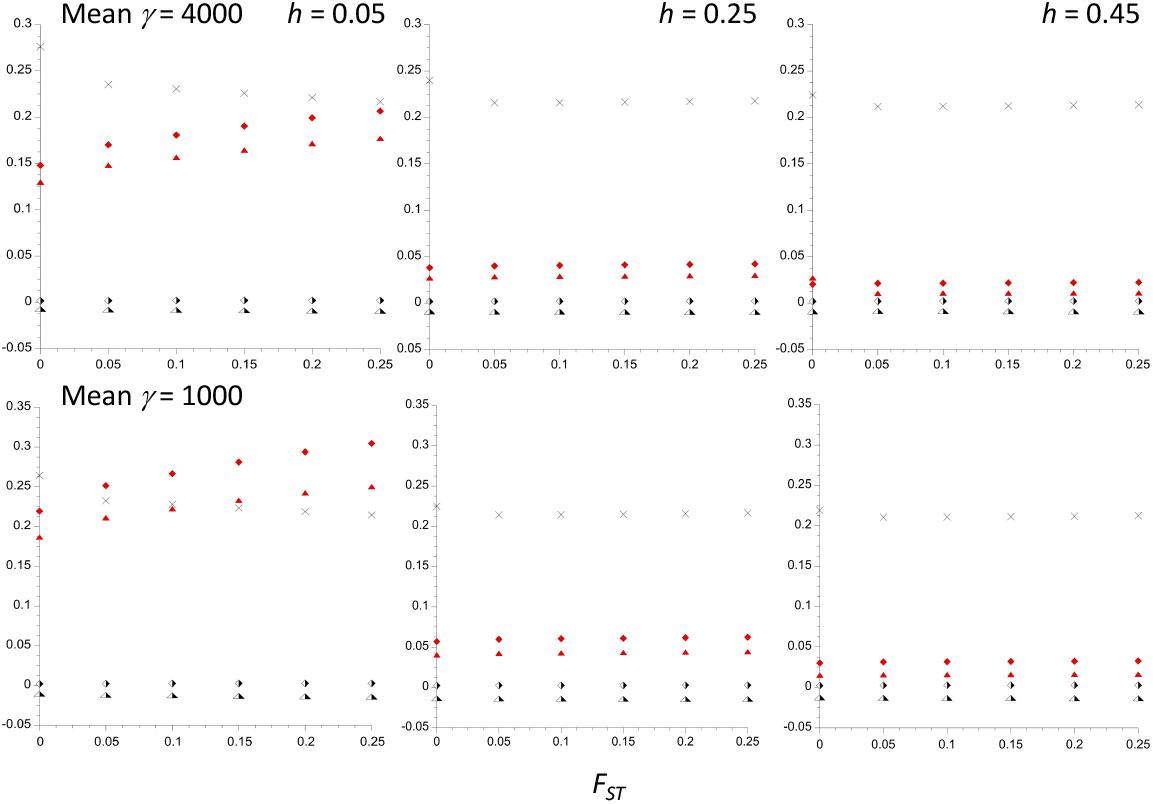
The mutational load and polymorphism statistics for an inversion polymorphism with inversion frequency *x* = 0.1 in a subdivided island population of total size *N*_*T*_ = 10^6^ with 200 demes and 10^5^ selected sites, plotted against *F*_*ST*_ for neutral sites independent of the inversion. The result for three different values of the dominance coefficient, *h*, are shown. The selection coefficients follow a gamma distribution with a mean of 5 x 10^−4^ and a shape parameter of 0.3. The mutation rate to deleterious alleles is 5 x 10^−9^ per basepair, with a mutational bias towards deleterious variants of 1.5. The upper and lower panels have products of 2*N*_*T*_ and mean *s* of 4000 and 1000, respectively. The filled and half-filled symbols denote values for the *In* and *St* subpopulations, respectively. The lozenges are the net mutational loads within the respective karyotypes, and the triangles are the selection coefficients against homokaryotypes relative to the heterokaryotype. The crosses are the ratios of the nucleotide site diversities for *In* versus the *St* subpopulations.

The relation with *F*_*ST*_ of the ratio of diversities at selected sites for *In* versus *St* with *x* = 0.1 is also shown in Figure 5, and is similarly rather weak. This result also applies to the population genomic statistics that are not displayed in Figure 5 − see Supplementary Table 4 for details (the Δ*θ*_*w*_ statistic was not calculated, as this statistic is hard to evaluate with population subdivision and was not very informative in the single population case). For example, with *x* = 0.1,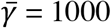 and *h* = 0.05, the ratio of *π*_*sel*_/*π*_*neut*_ for *In* versus *St* decreases from 2.32 with *F*_*ST*_ = 0 to 1.80 with *F*_*ST*_ = 0.25, indicatiing a greater efficacy of selection on the smaller *In* subpopulation when there is greater subdivision. With *h* = 0.25, the change is much smaller, from 1.97 to 1.90. Similarly, the ratio of the mean frequency of A_2_ for *In* versus *St* with *h* = 0.05 changes from 3.02 with *F*_*ST*_ = 0 to 3.78 with *F*_*ST*_ = 0.25, but only from 2.21 to 2.32 with *h* = 0.25.

These results are all for a relatively large deme size of *N* = 5000. Intuitively, it might seem that reducing the deme size would enhance the effects of drift within demes, and lead to larger loads and magnitudes of the selection coefficients, as well as reducing the values of such signatures of purifying selection as the mean frequencies of mutant alleles and the ratios of diversities at selected sites to neutral sites. The example in the upper part of Table 2, where deme sizes of 500 and 5000 are compared for the same neutral *F*_*ST*_ value, shows that this expectation is met for the case of equal frequencies of *In* and *St*, although the effects are small and are only visible in the table in a few cases for the mean fitnesses of the homokaryotypes and selection coefficients. However, the mean frequencies of mutations and the ratios of diversity at selected sites versus neutral sites show a clear pattern of reduced efficacy of selection with the smaller deme size, which is reduced in size by larger *h* and *F*_*ST*_. Further results for cases with smaller deme sizes are shown in Supplementary Table 5.

**Table 2.** Some load and population genomics statistics for a subdivided population of total size 10^6^ for two different local deme sizes (results for *N* = 500 and 5000 are shown in the upper and lower parts of each cell, respectively

| $x = 0.5$ | $\bar{w}_1$ | $t_1 = t_2$ | $\bar{q}_1 = \bar{q}_2$ | $\pi_{sel1}/\pi_{neut1}$ |
| --- | --- | --- | --- | --- |
| $F_{ST} = 0.05,$<br>$h = 0.05$ | 0.992<br>0.992 | 0.005<br>0.005 | 0.090<br>0.088 | 0.341<br>0.325 |
| $F_{ST} = 0.05,$<br>$h = 0.25$ | 0.994<br>0.998 | 0.002<br>0.002 | 0.087<br>0.085 | 0.262<br>0.254 |
| $F_{ST} = 0.05,$<br>$h = 0.45$ | 0.995<br>0.995 | 0.0003<br>0.0003 | 0.086<br>0.083 | 0.229<br>0.226 |
| $F_{ST} = 0.25,$<br>$h = 0.05$ | 0.987<br>0.987 | 0.090<br>0.083 | 0.101<br>0.099 | 0.263<br>0.258 |
| $F_{ST} = 0.25,$<br>$h = 0.25$ | 0.994<br>0.998 | 0.002<br>0.002 | 0.089<br>0.087 | 0.238<br>0.232 |
| $F_{ST} = 0.25,$<br>$h = 0.45$ | 0.995<br>0.995 | 0.0003<br>0.0003 | 0.086<br>0.084 | 0.228<br>0.221 |

| $x = 0.1$ | $\bar{w}_1/\bar{w}_2$ | $t_1$ | $t_2$ | $\bar{q}_1/\bar{q}_2$ | $(\pi_{sel1}/\pi_{neut1})$<br>$/(\pi_{sel2}/\pi_{neut2})$ |
| --- | --- | --- | --- | --- | --- |
| $F_{ST} = 0.05,$<br>$h = 0.05$ | 0.780<br>0.780 | 0.210<br>0.211 | -0.012<br>-0.012 | 3.23<br>3.29 | 2.17<br>2.04 |
| $F_{ST} = 0.05,$<br>$h = 0.25$ | 0.944<br>0.944 | 0.042<br>0.042 | -0.014<br>-0.014 | 2.26<br>2.29 | 1.93<br>1.88 |
| $F_{ST} = 0.05,$<br>$h = 0.45$ | 0.971<br>0.970 | 0.016<br>0.016 | -0.013<br>-0.013 | 1.93<br>1.96 | 1.88<br>1.85 |
| $F_{ST} = 0.25,$<br>$h = 0.05$ | 0.739<br>0.739 | 0.250<br>0.250 | -0.014<br>-0.014 | 3.72<br>3.78 | 1.83<br>1.88 |
| $F_{ST} = 0.25,$<br>$h = 0.25$ | 0.941<br>0.942 | 0.045<br>0.042 | -0.015<br>-0.015 | 2.29<br>2.32 | 1.91<br>1.90 |
| $F_{ST} = 0.25,$<br>$h = 0.45$ | 0.969<br>0.970 | 0.017<br>0.016 | -0.014<br>-0.014 | 1.93<br>1.96 | 1.90<br>1.86 |

$\bar{w}_1$ and $\bar{w}_2$ are the mean fitnesses of the products of random mating among *In* and *St* karyotypes, respectively.
The selection and mutation parameters are the same as in Figure 1.

The patterns are somewhat more complex, however, when there is a large difference in frequency between arrangements, as shown in Table 2 for the case of *x* = 1. Here, the differences in the load statistics between the large and small deme size cases are negligible, and probably cannot be distinguished from rounding errors. There is, however, a signature of a slightly enhanced efficacy of selection with the smaller deme size in the *In* subpopulation relative to *St*, indicated by a consistently smaller value of 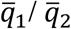 when *N* = 500. Confusingly, *π*_*sel*_/*π*_*neut*2_ is greater than *π*_*sel 2*_/*π*_*neut2*_ when *N* = 500, indicating the opposite pattern. The first of these results is explained by the fact that the much smaller size of the *In* subpopulation means that in both cases mutations are behaving nearly neutrally within demes, so that a lower deme size will not have much of an effect, whereas it will reduce the overall efficacy of selection on the *St* subpopulation, resulting in an increase in 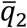. For example, with *h* = 0.05 and *F*_*ST*_ = 0.05, the ratio of 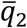 for *N* = 500 versus 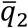 for *N* = 5000 is 1.024, whereas the value for 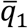 is 1.006. The corresponding ratio for *π*_*sel*_/*π*_*neut*1_ is 1.072 whereas that for *π*_*sel 2*_/*π*_*neut2*_ is 1.005. This is entirely due to a higher ratio of *π*_*sel*1_ for *N* = 500 versus *N* = 5000, since there is no effect of local deme size on neutral diversity for a fixed neutral *F*_*ST*_. It is not entirely clear how to interpret this pattern, but one factor is likely to be the fact that the relaxation of the efficacy of selection on *St* leads to an increase in *F*_*ST*_ among demes at selected sites, which would work against any increase in *π*_*sel 2*_ due to a reduced efficacy of selection.

## Discussion

It should be borne in mind that the results described above relate to a situation in which an inversion polymorphism is maintained by balancing selection that is invariant over space and sufficiently strong that the frequency of the arrangements is approximately constant over time and space. This obviously does not apply to many natural situations, e.g., when there are clinal patterns of variation in inversion frequencies, as is often the case (Krimbas and Powell 1992). Nevertheless, the theoretical results described above address several questions that can in principle be answered by comparisons with empirical studies. First, does a low frequency arrangement accumulate a larger mutational load than its counterpart? Second, can mutational load contribute a significant selective advantage to heterokaryotypes, which might help to stabilize the polymorphism? Third, can differences in population genomic statistics between common and rare arrangements shed light on differences in mutational load between arrangements? These questions are each discussed in turn below.

### Are the relative values of the fitness components of carriers of different gene arrangements consistent with mutational load?

The relevant variables with respect to the relative fitnesses of the *In* and *St* subpopulations are the *L*_*i*_, the mutational loads associated with individuals produced by random mating among outbred individuals homozygous for karyotype *i*, where *i* = 1 for the rarer arrangement (*In*) and *i* = 2 for the more common one (*St*). These are, course not directly observable, since the fitness of mutation-free individuals is unknown, but they can be used to predict the relative values of the mean fitnesses (or fitness components) for *In* and *St*. On the assumption of multiplicative fitnesses, the ratio of the mean fitnesses of type 1 and type 2 homokaryotypes is 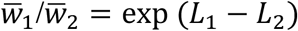, corresponding to a difference *L*_1_ − *L*_2_ in the natural logarithm of mean fitness. The mutational and selection parameters used here (a mutation rate of 5 x 10^−9^ per site and a gamma distribution of selection coefficients with shape parameter 0.3) are consistent with the results of population genomic studies of *D. melanogaster*, e.g., Kousathanas and Keightley (2013) and Assaf et al. (2017).

In order to relate the theoretical predictions to data on inversion polymorphisms, it is necessary to have information about *h* and the parameters of the distribution of *s*, which is made difficult by the fact that there is much uncertainty about the relation between *h* and *s*. The evidence from *Drosophila* studies of the effects of mutations on fitness components suggest that only very strongly selected deleterious mutations such as homozygous lethals have *h* values low as 0.05, whereas the much more abundant deleterious mutations with *s* ≤ 0.02 have *h* values of the order of 0.25 or more (Crow 1993; Manna et al. 2011). It therefore seems safe to use *h* = 0.25 as a working value for comparing theory with data, since mutations like lethals that are strongly selected against when heterozygous will be held close to their deterministic equilibrium values in both the *In* and *St* subpopulations (Nei 1968), and hence will not contribute to the genetic differences between them. It is worth noting that a higher load for the rarer arrangement does not require mutations to have *h* < ½, contrary to what is often stated (e.g., Jay et al. 2021). If drift is sufficiently strong in relation to selection that there is a higher expected frequency of deleterious mutations in the smaller subpopulation, a higher load can arise even for completely dominant mutations.

Estimates of the distribution of *γ* = 2*N*_*e*_*s* from population genomics studies reflect the more weakly selected part of the distribution of selection coefficients and are the most useful source of information for the present purpose. With a shape parameter of 0.3 and *h* = 0.25, *π*_*sel*2_/*π*_*neut*2_ = 0.15 with 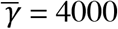 when *x* = 0.1. This is only slightly higher than the observed ratios of nonsynonymous to silent site diversities in normally recombining regions of the genome in putatively ancestral range populations of *D. melanogaster* (e.g., Campos et al. 2014), suggesting that this value of 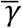 can used as a working estimate for nonsynonymous mutations.

One of the best-studied *D. melanogaster* inversions is *In(3R)P*, with a frequency of 0.1 in populations in its ancestral range in Africa (Kapun and Flatt 2019), which are the most relevant populations for comparisons with the theoretical predictions. It is in the middle of the size range for polymorphic inversions in this species and covers approximately 8 Mb of sequence (Kapun et al. 2023); This corresponds to approximately 1000 protein coding sequences, i.e., 10^6^ nonsynonymous sites rather than the 10^5^ selected sites illustrated in the figures. Functional noncoding sequences may also contribute to the mutational load, and these appear to be under weaker selective constraints than nonsynonymous mutations (Andolfatto 2005; Casillas et al. 2007; Campos et al. 2017). As a rough estimate of their contribution to the load and population genomic statistics, it is plausible to assume that functional non-coding sites are three times as abundant as nonsynonymous sites (Halligan and Keightley 2006) but have 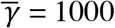 rather than 4000. With *x* = 0.1 and *h* = 0.25, the results in Supplementary Table 2 imply that, by combining the effects of 10^6^ nonsynonymous sites with 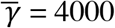 and 3 x 10^6^ noncoding sites with 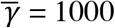 and *h* = 0.25, we would have *L*_1_ = 2.09 and *L*_2_ = 0.08 for an inversion with frequency 0.1, giving a predicted 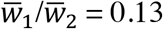.

Predictions of this kind are highly sensitive to the frequency of the inversion, and to assumptions concerning the abundance of functional sites and the strength and mode of selection. With the model just described and an inversion frequency of 0.3, we have *L*_1_ = 0.42 and *L*_2_ = 0.11, giving 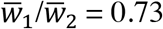. If only nonsynonymous sites are considered, 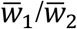 rises to 0.69 for *x* = 0.1 and 0.95 for *x* = 0.3. Synergistic epistasis among deleterious mutations is expected to reduce the *L*_*i*_ by a factor of approximately two (Kondrashov 1995; Charlesworth 2013), so that the model of a mixture of 10^6^ nonsynonymous and 3 x 10^6^ functional non-coding sites would generate 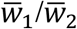 values of 0.37 and 0.86 for *x* = 0.1 and *x* = 0.3, respectively. These complexities means that it is difficult to make rigorous comparisons between theory and data, and the estimate of 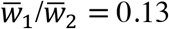 for *In(3R)P* is probably at the lower end of what is expected.

A large number of studies of the effects of inversions on fitness and components of fitness under laboratory conditions have been published, with *D. melanogaster* and *D. pseudoobscura* being been the most intensively studied species − see reviews by Sperlich and Pfriem (1986), Krimbas and Powell (1992) and Kapun and Flatt (2019). Many of these studies do not, however, provide useful information for the present purpose, either because they involve traits like male mating success that are hard to quantify in terms of relative fitnesses of the type used here, or because only small numbers of independently sampled arrangement haplotypes were used, or because samples from laboratory rather than natural populations were used. The most informative available estimates are shown in Supplementary File S4, with their standard errors where available.

For the two large-scale studies of egg-to adult viability in *D. melanogaster* (Mukai and Yamaguchi 1974; Watanabe et al. 1976), where balancer crosses were used to extract 2^nd^ or 3^rd^ chromosomes from wild flies, the crosses involving pairs of independently extracted chromosomes were divided into cases where either each member of a pair was inversion-free or at least one member of each pair carried an inverted chromosome. There is no evidence for a difference between these categories in either study. One possible explanation for this discrepancy is the fact that a single component of fitness such as viability reflects only a portion of the net effect of deleterious mutations on fitness. Table S8 in Charlesworth (2015) presents estimates of the ratios of the fitness effects of deleterious mutation on various fitness components to their effects on net fitness (the *α* parameters). The estimates of *α* are subject to considerable uncertainty, but even the lowest estimate for viability (0.09) cannot explain the absence of effects on viability in these experiments.

Studies such as these that involve populations of *D. melanogaster* outside the ancestral range of the species in south-eastern Africa are difficult to interpret, because of the effects on variability of the severe population bottlenecks associated with the spread of the species out of Africa (e.g., Haddrill et al. 2005). Indeed, a curious feature of the data of Mukai et al. (1974) on the 2^nd^ chromosome is the much higher frequency of homozygous lethal *In* chromosomes than *St* chromosomes (54% versus 36%); this is associated with a higher frequency of pairs of crosses in which combinations of different lethal-bearing chromosomes were lethal (19% versus 11%), suggesting that there may have been a recent population size bottleneck that affected the *In* chromosomes more severely than the more abundant *St* chromosomes. Unfortunately, no information on the fitness effects of inversion genotypes appears to be available for samples from the ancestral range of *D. melanogaster*.

Species that are less subject to this problem, such as *D. pseudoobscura*, are thus more favorable material. Estimates of the net relative fitnesses and of two measures of viability for *D. pseudoobscura* 3^rd^ chromosome arrangements in population cages are also given in Supplementary File S4 (the horizontal lines separate data from different experiments). AR and CH cover approximately 30% and 20% of chromosome 3 (the homolog of chromosome arm 2R of *D. melanogaster*), respectively (Powell 1992), and are thus suitable for comparison with the theoretical prediction for *In(3R)P* given above. For the viability experiments of Dobzhansky (1947), involving CH and ST from a California population, CH had a frequency of between 0.20 and 0.35 in the original population, but which varied substantially over the year (Wright and Dobzhansky 1946, Fig. 2). The viability of CH/CH individuals was 86% of that of ST/ST individuals, which is reasonably consistent with expectations for an inversion with a frequency of around 0.3 The relative viabilities of CH/CH and AR/AR individuals were similar, as expected from the similar frequencies of CH and AR in this population.

The results for egg-to-adult viability measurements that used a balancer chromosome to extract 3^rd^ chromosomes from a natural population of *D. pseudoobscura* (Crumpacker and Salceda 1968) show smaller effects than those predicted from mutational load; only the results for the two most common arrangements are shown in the table, with AR and CH having frequencies of approximately 0.5 and 0.28 in the population; the ratio of viabilities of CH/CH versus AR/AR in crosses between carriers of independently extracted chromosomes is 0.98, which is unlikely to differ significantly from 1. The net fitness estimates obtained from population cage experiments on the natural population used for Dobzhansky’s viability estimates, but which segregated for ST, AR and CH (Wright and Dobzhansky 1946), show patterns that are inconsistent with the mutational load predictions; in particular, AR/AR has a much lower fitness than CH/CH despite their similar frequencies. In contrast, the ratios of net female fitnesses for CH/CH to AR/AR estimated by Anderson and Watanabe (1997) for a laboratory population derived from the same population but segregating only for CH and AR is 0.85, which is consistent with the fact that the population equilibrated at about 25% CH. However, the ratio for ST/ST versus AR/AR in another experimental population was 0.70, despite the fact that ST is usually at least as frequent as AR (Powell 1992).

The overall conclusion from these analyses of the relative performance of rare versus common arrangements is that some measurements fit the expectation of a larger equilibrium mutational load for the less frequent *Drosophila* inversions, but the overall patterns strongly suggest that other factors obscure the contribution of load to homokaryotype fitnesses.

### Can mutational load create a selective advantage to heterokaryotypes?

The theoretical results described earlier make it clear that the mutational load model used here can only create a heterokaryotypic advantage when *In* and *St* are present at nearly equal frequencies (see Figures 1 and 2, and Supplementary Tables 2 and 3); furthermore, the magnitude of such an advantage is likely to be small. For example, with *x* = 0.5, *h* = 0.25 and 10^5^ selected sites, the selection coefficient against both homokaryotypes relative to the heterokaryotype given by Equations (3) is 0.0016, with 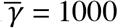 and 0.0010 with 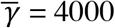. However, if the above model of 3 x 10^6^ sites with 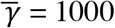 and 10^6^ sites with 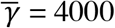 is used, the selection coefficient becomes 0.057, which is small but non-negligible. With *x* = 0.4, the selection coefficients under this model are for *t*_1_ = 0.13 for *In*/*In* and *t*_2_ = 0.0013 for *St*/*St*. For lower values of *x*, the selective advantage to *In*/*St* heterokaryotypes over *St*/*St* is replaced by a selective disadvantage, as shown in Figures 1 and 2.

The reason for this behavior is that a rarer arrangement accumulates a larger mutational load than its counterpart, due to the lower efficacy of selection with smaller *N*_*e*_ (see the Introduction). When a haplotype from the *In* population is made heterozygous with a haplotype from the *St* subpopulation, there is a smaller expected number of heterozygous mutations in the *In*/*St* individuals than in the *In* subpopulation. This means that *t*_1_ > 0 if *h* < ½, as is evident from Equation (3a), where *F*_1_, 1 − *F*_1_, and *C*_12_ are all positive, as is the difference <*δq*> between the *In* and *St* subpopulations in the expected frequency of mutations. In contrast, *In*/*St* individuals have a larger expected number of mutations than the *St* subpopulation. As can be seen from Equation (3b), if *h* < ½ and the magnitude of <*δq*> is sufficiently large, *t*_2_ is negative. But if the two arrangements are equally frequent, <*δq*> = 0, and the remaining terms guarantee that *t*_1_ = *t*_2_ > 0 when *h* < ½. In all cases, *t*_1_ = *t*_2_ = 0 if *h* = ½. The only surprising aspect of the theoretical results is that *t*_2_ is so sensitive to the effect of the relative subpopulation sizes on <*δq*>.

These theoretical predictions can be compared with the data in Supplementary File S4. The two studies of viability in *D. melanogaster* showed no difference in viability between chromosomal heterozygotes that were free of inversions and chromosomal heterozygotes where at least one of the pairs of chromosomes involved carried an inversion. Since the inversions were rare, most of the latter cases will have involved heterokaryotypes, so the lack of any difference is consistent with an absence of heterokaryotypic superiority, as expected for rare inversions. For *D. pseudoobscura*, the results on net fitness and on viability from population cage experiments indicate strong heterokaryotypic advantages, much larger than the theoretical predictions for inversions of the size involved here. In contrast, the balancer cross data on viability showed a small heterokaryotypic advantage (Crumpacker and Salceda 1968), consistent with the theoretical predictions for equally frequent arrangements. Mérot et al. (2020), Huang et al. (2022) and Pei et al. (2022) found no evidence for heterokaryotypic superiority for several fitness components in seaweed flies, sunflowers and zebra finches, respectively. Overall, it seems likely that the mutational load model may contribute modestly to heterokaryotypic superiority for inversions that are at intermediate frequencies, but cannot explain the large net fitness effects seen in the *D. pseudoobscura* inversions. This parallels the finding that mutational load is unlikely to provide a selective advantage to new autosomal inversions in randomly mating populations (Nei et al. 1967; Connallon and Olito 2021; Jay et al. 2022).

### Population genomic indicators of a reduced efficacy of selection on low frequency arrangements

As described in the section *A single population: population genomics statistics*, the two most informative population genomic statistics concerning a reduced efficacy of selection on a low frequency arrangement are the mean frequencies of variants at the selected sites (the 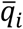) and the ratios of diversities at selected and neutral sites (the *π*_*seli*_/*π*_*neuti*_). The measure of skew towards low frequency variants relative to neutral expectation (Δ*θ*_w*i*_) is much less sensitive to subpopulation size, unless the dominance coefficient is implausibly small. In addition, measures of skew are sensitive to population size changes (Tajima 1989), and must be treated with caution when making inferences about selection. For this reason, only the other two statistics will be considered here.

A problem with the use of the 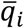 is that these are not directly observable unless bioinformatic methods for inferring variants as deleterious are used; simply using the mean frequency of derived nonsynonymous variants in a sample as a proxy (cf., Campos, et al. 2014) is not necessarily adequate when selection is weak, since it does not take into account fixed sites. Stenløk et al. (2022) used PROVEAN scores to estimate the mean numbers of deleterious missense mutations in inverted and standard arrangements of Atlantic salmon, but found no significant differences; an enrichment of small indels in the large (3.09 Mb) Chr18 inversion was, however, detected. The frequencies of this inversion are, however, variable between populations, so it is not clear how to interpret this difference.

There are, however, theoretical reasons for expecting the ratio 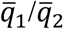 to behave similarly to the ratio *R* _*π*_ = (*π*_*sel*1_/*π*_*neut*2_)/ (*π*_*sel*2_/*π*_*neut*2_), at least when *h* is not too small. Welch et al. (2008) analyzed the properties of the ratio of selected site to neutral site diversity under a similar model to that used here, assuming *h* = 0.5 and a gamma distribution of selection coefficients. They showed that with 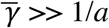, where *a* is the shape parameter of the gamma distribution in Equation (9), the following approximate relation holds:

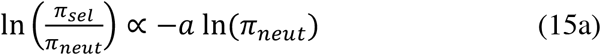

In the present case, this implies that:

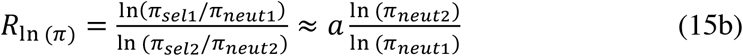

While we would not expect this relation to be exact for *h* < 0.5, it is plausible to assume that it would be a reasonably good approximation when *h* is not too close to 0. The argument used by Welch et al. (2008) implies that a similar relation should apply to the mean frequencies of deleterious mutations:

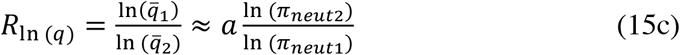

The accuracy of these approximations can be tested using the numerical results in Supplementary Tables 2 and 3. Table 3 gives some examples for the case of a single randomly mating population, showing that the two ratios on the left-hand sides of Equations (15b) and (15c) behave very similarly as functions of *h*, with 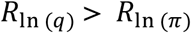 when *h* < 0.25, approaching the prediction on the right-hand sides of the equations for *h* ≥ 0.35. Similar results apply to the case of a subdivided population.

**Table 3.** Values of the ratios with respect to *In* versus *St* of the natural logarithms of *π* _*sel*_/ *π* _*neut*_ (*R*_*ln(π*_) and 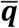 (*R*_*ln(q)*_) for the case of a single population

| $h$ | $\bar{\gamma} = 1000$ | | | | $\bar{\gamma} = 4000$ | | | |
| --- | --- | --- | --- | --- | --- | --- | --- | --- |
| | $x = 0.1$ | | $x = 0.3$ | | $x = 0.1$ | | $x = 0.3$ | |
| | $R_{ln(\pi)}$ | $R_{ln(q)}$ | $R_{ln(\pi)}$ | $R_{ln(q)}$ | $R_{ln(\pi)}$ | $R_{ln(q)}$ | $R_{ln(\pi)}$ | $R_{ln(q)}$ |
| 0.05 | 0.840 | 1.106 | 0.395 | 0.485 | 0.884 | 1.110 | 0.403 | 0.487 |
| 0.15 | 0.720 | 0.907 | 0.313 | 0.393 | 0.744 | 0.909 | 0.319 | 0.394 |
| 0.25 | 0.679 | 0.794 | 0.282 | 0.329 | 0.694 | 0.795 | 0.286 | 0.330 |
| 0.35 | 0.661 | 0.718 | 0.267 | 0.285 | 0.675 | 0.719 | 0.270 | 0.285 |
| 0.45 | 0.653 | 0.658 | 0.257 | 0.250 | 0.665 | 0.659 | 0.261 | 0.250 |
| 0.50 | 0.650 | 0.632 | 0.254 | 0.235 | 0.662 | 0.633 | 0.257 | 0.235 |

The predicted values of the ratios with the shape parameter $\alpha = 0.3$ are 0.659 for $x = 0.1$ and 0.254 for $x = 0.3$ .
The mutational parameters and population size are the same as in Figure 1.

These results suggest that *R*_*π*_ can used as a conservative proxy for 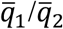, at least for the case of a gamma distribution of selection coefficients with 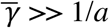 and an intermediate dominance coefficient. An objection to using the correlation between 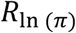 and a measure of neutral diversity such as synonymous site diversity to investigate whether the efficacy of purifying selection declines with *N*_*e*_ is that *π*_5*neut*_ is the denominator of *π*_*sel*_ /*π*_*neut*_. In addition to the statistical problem of a negative correlation introduced by this relationship, discussed by James et al. (2017), it could be argued that sites under sufficiently strong purifying selection would maintain a constant diversity across different *N*_*e*_ values (Campos et al. 2014). If this were the case, the expectation of *π*_*sel*_/*π*_*neut*_ would simply be proportional to 1/*π*_*neut*_, and we would then have *R*_*π*_ = *π*_*neut2*_/*π*_*neut*_. As described in the *Results* section *A single population: population genomics statistics*, this is not the case; *R* _*π*_ with *x* < 0.5 is always less than *y*/*x*. For example, with 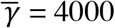 and *h* = 0.25, *R*_*π*_ = 2.00 for *x* = 0.1 (*y*/*x* = 9) and *R*_*π*_ = 1.33 for *x* = 0.3 (*y*/*x* = 2.33). This indicates that *R*_*π*_ provides a signal that purifying selection is weakened by smaller subpopulation size. Comparisons of this kind could easily be done using real data.

Overall, therefore, these considerations suggest that, despite the above reservations, *R* _*π*_ is quite a useful index of the efficacy of purifying selection, and that one might expect 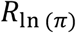 to be approximately equal to *a* ln(*π*_*neut*2_)/ ln(*π*_*neut*_). A related principle was used by James et al. (2017) and Castellano et al. (2018) to test for relations between the efficacy of purifying selection and *N*_*e*_ for animal mitochondrial genomes and different regions of the *D. melanogaster* nuclear genome, respectively. Unfortunately, there appears to be relatively little relevant information for autosomal inversions, other than cases such as the mimicry supergene in *Heliconius numata* (Jay et al. 2021) and the behavioral supergene of the white-throated sparrow *Zonotrichia albicolis* (Jeong et al. 2021), which are largely maintained as heterozygotes due to negative assortative mating. These systems are thus analogous to sex chromosomes, where one arrangement is permanently heterozygous and effectively lacks recombination. There is thus likely to be intense Hill-Robertson interference (Charlesworth and Charlesworth 2000), which would greatly reduce the efficacy of selection below the simple effect of a lower subpopulation size. This is consistent with the strongly elevated *π*_*sel*_/*π*_*neut*_ values found for the *H. numata* inversions; *Z. albicolis* showed only a modest effect. Jay et al. (2021) also found a large increase in the density of transposable elements (TEs) in the *H. numata* inversions and interpreted this as evidence for an increased mutational load. The accumulation of TEs in low recombination regions of genomes, including low frequency *Drosophila* inversions (Sniegowski and Charlesworth 1994), has long been documented (Charlesworth et al. 1994). Most insertions are found in intergenic regions, where direct selective effects are likely to be weak, and where ectopic exchange inducing deleterious chromosome rearrangements is probably a major factor in causing their elimination. It is thus likely that a reduced frequency of ectopic recombination is the major factor in causing higher densities of TE insertions in such cases (Charlesworth et al. 1994), so that this phenomenon cannot be taken as evidence for an increased mutational load.

### What strength of selection has the main effect on the load and population genomic statistics?

Another question raised by the theoretical results is: what part of the distribution of selection coefficients contribute to the differences between the *In* and *St* subpopulations? Stronger selection reduces the frequencies of deleterious mutations and their chances of fixation within a subpopulation, but also increases the sizes of any resulting loads. The major contribution to the relevant load statistics is thus likely to come from selection coefficients that are neither too large nor too small. This expectation can be tested by examining the contributions from the different zones described in the section *A single population: obtaining the mutational load and population genomic statistics*, which are shown in Supplementary Table 2 for the case of a single population. These results show that the major contributions to the *L*_*i*_ and *t*_*i*_ come from zone 2a, defined by an intermediate intensity of selection (for more details, see Supplementary Table 6). For example, for a randomly mating single population with 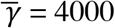, *x* = 0.1 and *h* = 0.25, zones 1, 2a, 2b and 3 contribute 4%, 36%, 24% and 36%, respectively, to the distribution of selection coefficients against deleterious mutations. Zone 2a alone gives *L*_1_ = 0.0375, *L*_2_ = 0.0012, *t*_1_ = 0.0270 and *t*_2_ = − 0.0090, compared with net values of *L*_1_ = 0.0381, *L*_2_ = 0.0016, *t*_1_ = 0.0271 and *t*_2_ = − 0.0090. It covers the interval (0.278, 463) of *γ* for the whole population.

The finer dissection of the distribution of selection coefficients used in the case of a subdivided population reveals that the so-called quasi-neutral zone 2 in this case (see the Appendix) contributes most to the load statistics (Supplementary Table 3). For example, with 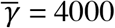, *x* = 0.1, *h* = 0.25 and *F*_*ST*_ = 0.05 (*M* = 19), zones 1, 2, 3 and 4 for the subdivided case contribute 4%, 17%, 21% and 58%, respectively. Zone 2 covers the *γ* interval (0.278, 55.6), and contributes *L*_1_ = 0.0344, *L*_2_ = 0.0011, *t*_1_ = 0.0249 and *t*_2_ = − 0.0082, compared with values of *L*_1_ = 0.0401, *L*_2_ = 0.0019, *t*_1_ = 0.0285 and *t*_2_ = − 0.0094 for the whole distribution. In this case, with 200 demes of size 5000 each, both the *In* and *St* subpopulations are behaving effectively neutrally within demes for this part of the distribution of selection coefficient. A similar pattern holds even with *F*_*ST*_ = 0.25 (*M* = 3). Nonetheless, the results are very similar to those for the single population, implying that the expectations of the population genetic parameters for an island model with a relatively low level of isolation between demes are mainly controlled by the size of the metapopulation, as was shown to be the case analytically by Wakeley (2003). Intuitively, this reflects the fact that, under these conditions, a mutation spends relatively little of its total sojourn time in the deme in which it arose.

### Confounding factors

The present study assumes that selected sites are at statistical equilibrium under mutation, selection and drift, an absence of recombinational exchange between *In* and *St* in heterokaryotypes, and complete independence among sites under selection. These assumptions are likely to violated in many real-life situations. First, consider the question of departure from equilibrium. If the inversion arises as a unique mutational event, as the available evidence seems to suggest (reviewed in Charlesworth 2023), the *In* subpopulation will initially completely lack genetic variability, and much time will be needed for equilibrium to be approached. The *St* subpopulation can be assumed to be close to equilibrium initially and will thus approach its new equilibrium much faster than the *In* subpopulation, so that only the latter need be considered here. For this subpopulation, the magnitudes of the *L*_*i*_, *B*_*i*_ and *t*_*i*_ will be below their equilibrium values for a long time after the inversion has approached its equilibrium frequency under balancing selection.

It is difficult to make exact predictions about the rate of approach to statistical equilibrium when both drift and selection play a role, which has been shown above to be the situation that contributes the most to the selective differences among karyotypes. For the limiting case of complete neutrality, it is known that the divergence of the expected nucleotide site diversity from its equilibrium value at time *t* is equal to the product of its initial value and exp (-*t*/2*N*_*e*_) in a randomly mating population (Malécot 1969, p.40), so that the timescale for approach to equilibrium is of the order of 2*N*_*e*_ generations. For the other limiting case of fully deterministic evolution, with *γ* >> 1, the divergence at time *t* of the mutant allele frequency *q* from its equilibrium value of *u*/*hs* is approximately equal to the product of its initial value and exp (-*t*/*hs*). For the intermediate situation when *γ* is not much greater than one, these two measures of the rate of approach to equilibrium are not very different, so it plausible to assume that the true rate lies between them. The time needed to approach equilibrium with respect to mutations that have the largest effect on the load statistics is thus likely to be < 2*Nx* generations under a Wright-Fisher model. For a population with an *N*_*e*_ of 10^6^ and *x* = 0.1, this would correspond to 10^5^ generations, i.e., about 10,000 years for a species like *D. melanogaster* with approximately 10 generations per year. For *x* = 0.5, about 50,000 years would be required.

A problem with assessing whether these timescales are consistent with data on inversion polymorphisms is that recombinational exchange between *In* and *St* due to gene conversion, for which there is much evidence in *Drosophila* (Korunes and Noor 2019), means that estimates of inversion age based on sequence divergence between different arrangements tend to produce underestimates of age (Charlesworth 2023). A variety of lines of evidence suggests, however, that inversions such as *In(3R)P* are close to selection-drift-recombination equilibrium with respect to neutral variability (Charlesworth 2023); since selection against deleterious mutations will cause a faster approach to equilibrium than with neutrality, it is likely that the load statistics will also be close to their equilibrium values, unless demographic disturbances have caused serious perturbations.

However, the theory developed here has ignored recombination. If the estimate of a typical rate of gene conversion of 10^−5^ in female meiosis in inversion heterokaryotypes in *Drosophila* (Korunes and Noor 2019) is accepted, the effective rate is 0.5 x 10^−5^ due to the absence of exchange in males. An *hs* value that somewhat exceeds 10^−5^ would thus be sufficient to overcome the effects of gene flow between *In* and *St*; with *h* = 0.25 and *N*_*T*_ = 10^6^, this would correspond to *γ* > 80, which lies outside the range of *γ* values that contribute to a noticeable difference in load statistics between *In* and *St* with *x* = 0.1, as discussed in the previous section. It is therefore likely that recombination will significantly reduce such differences, providing another reason for regarding the above estimates as providing upper bounds to the predictions.

In addition, the fitness effects of any deleterious mutations that were present on the initial inversion haplotype have apparently been ignored; however, the final equilibrium state considered here, where reverse mutations have been included, means that such effects will have been removed. During the approach to equilibrium they must, of course, play a role in reducing any selective advantage to a new inversion (Nei et al. 1967; Connallon and Olito 2021; Jay et al. 2022).

The low frequency of recombination in inversion heterokaryotypes for a rare inversion may create Hill-Robertson interference effects, as noted above in the section *Population genomic indicators of a reduced efficacy of selection on low frequency arrangements*. Berdan et al (2021) used simulations to study this possibility, and found strong interference effects under the assumption that mutations were completely recessive. As pointed out previously, this assumption is implausible, even for mutations with large homozygous fitness effects. In addition, recombination between *In* and *St* will reduce interference effects. These will tend to increase the load in the less frequent subpopulation and in heterokaryotypes, and so will not contribute to the maintenance of the inversion polymorphism. There is little evidence for any effects of Hill-Robertson interference on population genomic statistics for the *D. melanogaster* inversions (Charlesworth 2023).

## Conclusions

The theoretical results described here show that a long-maintained autosomal inversion polymorphism with no recombination in heterokaryotypes may develop a substantially higher mutational load for the less frequent arrangement. The magnitude of the difference between arrangements can be large for rare polymorphic inversions of the size usually encountered in *Drosophila* populations, but declines quickly as the frequency of the rare arrangement increases. It is also strongly influenced by the abundance of relative weakly selected non-coding sequences, since drift acts more strongly on these than on strongly selected nonsynonymous mutations. A selective advantage to heterokaryotypes is only expected when the two alternative arrangements are nearly equal in frequency, and is likely to be small even in this case. Experiments on the effects of several *Drosophila* inversion polymorphisms on fitness components give inconsistent results, although mutational load may contribute to some of the effects that have been detected. It should also be possible to detect molecular signatures of an increased load, such as an enhanced ratio of nonsynonymous to synonymous nucleotide site diversities, but the data are currently too scanty to draw firm conclusions. The effects of recombinational exchange in heterokaryotypes and Hill-Robertson interference, which oppose each other, were ignored here, and deserve further study.

## Data availability

No new data or reagents were generated for this work. The codes for the computer programs used to generate the results described above are available in Supplementary Files S2 and S3.

### Acknowledgments

I thank Deborah Charlesworth, Tim Connallon, Thomas Flatt and Michael Whitlock for their helpful comments on a draft of this paper.

## Appendix

### 1. Load calculations for a single population

Zone 1 corresponds to a quasi-neutral region where *γ* is sufficiently small (e.g., 0.25) that the selection term in Equations (5) can be ignored. The two *q*_*i*_ can be then treated as independent variables, each following a beta distribution with parameters *α*_*i*_ and *β*_*i*_. In this case, we have:

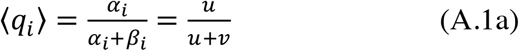

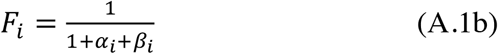

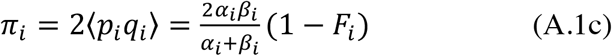

where Equation (A.1c) give the expected nucleotide site diversity for subpopulation *i*. In most applications, we have *α*_*i*_, *β*_*i*_ << 1, so that this equation can be approximated by:

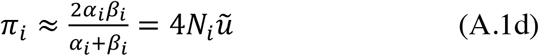

where ũ = 2*kv*/(1 + *κ*) is the net mutation rate at base composition equilibrium (Charlesworth and Charlesworth 2010, p.274). This is identical to the infinite sites model formula for nucleotide site diversity, which does not explicitly take reverse mutations into account (Kimura 1971).

These expressions can be inserted into Equations (1). For this zone, the exponential term in *γ* in the gamma distribution p.d.f. can be neglected. As it is assumed that *x* ≤ 0.5, the focus is on ensuring that the *St* population behaves as neutral. As shown in Section 3 of Supplementary File S1, if the upper bound to *γy* (the scaled selection coefficient for *St* in zone 1) is denoted by *γ*_*c*1_, the probability *P*_*z*1_ that *γy* ≤ *γ*_*c*1_ and the integral of *γy* over this zone, *I*_1_ are given by:

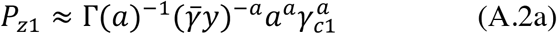

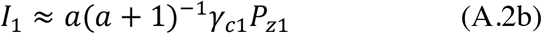

Zone 2 involves a moderate intensity of selection, with *γ*_*c*1_ ≤ *γy* ≤ *γ*_*c*2_, where *γ*_*c*2_ is such that there is a non-trivial probability that either *q*_1_ or *q*_2_ reaches an intermediate or high frequency. In order to increase the accuracy of the integrations of the joint p.d.f. over the distribution of *γ*, Zone 2 was divided into two subzones, with Zone 2a having an upper limit of *γy* = *γ*_*c*2*a*_, and Zone 2b with *γ*_*c*2*a*_ ≤ *γy* ≤ *γ*_*c*2_. Numerical values for *γ*_*c*1_, *γ*_*c*2*a*_ and *γ*_*c*2_ can be found in Supplementary Table 1. The probability of *s* falling into this zone, *P*_*z*2*a*_, is given by the excess over *P*_*z*1_ of the integral of the gamma distribution from 0 to *γ*_*c*2*a*_; the probability for zone 2b, *P*_*z*2*b*_ is found in a similar way by using *γ*_*c*2*a*_ and *γ*_c2_ as integration limits. The bivariate distribution of Equation (8) is used to determine the expected value of the load statistics in Equations (1) for a given *γ*, using the means and variances of the *q*_*i*_ generated by the distribution. The integrals over *γ*_c1a_ ≤ *γy* ≤ *γ*_c2a_ and *γ*_c2a_ ≤ *γy* ≤ *γ*_c2_ of the products of Equations (1a) − (1c) with *n*_*s*_ and the p.d.f. for the gamma distribution of *γy* yield the net contributions from Zone 2a and 2b to the load statistics.

It was found that use of Equation (8) for values of *γy* that were close to *γ*_*c*1_ (usually set to 0.25) gave inaccurate results, with expected frequencies of A_2_ somewhat greater than those obtained for Zone 1. To avoid this problem, an approximation to Equation (8), described in section 4 of Supplementary File S1, was used for values of *γy* less than a threshold value. Trial and error showed that accurate results, where both methods agreed closely in the results for integration over the whole zone were obtained by setting this threshold to 0.25*γy*/(*hx*), equivalent to 40*γy* with *h* = 0.05 and *x* = 0.5 (this adjusts for the fact that selection against rare mutations is effectively weaker when they are more recessive).

Zone 3 involves a high intensity of selection, such that *q*_1_ and *q*_2_ are both confined to their boundaries close to zero; here *γ*_*c*2_ ≤ *γy* ≤ *γ*_*c*3_, where *γ*_*c*3_ is assigned as the upper limit to the distribution of *γy*, usually chosen to correspond to the upper 99^th^ percentile of the gamma distribution. The probability of *s* falling into this zone, *P*_*z*3_, can be found as *P*_*z*2*a*_ for and *P*_*z*2*b*_. The two frequencies *q*_1_ and *q*_2_ can be treated as independent variables, since their expected product is negligible and A_2_ alleles within *In* and *St* haplotypes have little chance of encountering each other. Provided that *h* is sufficiently greater than zero, their p.d.f.s for a given *h* and *s* are well approximated by gamma distributions (Nei 1968), with shape parameters *α*_*i*_ and means *u*/*hs*. The corresponding variances are (*u*/*hs*)^2^/*α*_*i*_, allowing the *F*_*i*_ and other load statistics of Equations (1) to be determined easily. The net contributions of zone 3 to the load statistics can then be found by integrations of the same type as described for zone 2.

The integrals over the distribution of *γ* for each of Zones 2a, 2b and 3 were evaluated numerically using Simpson’s rule (Atkinson 1989), usually using 650 points over a single interval for Zones 2a, 2b and 3 (trial and error showed that this number was sufficient to produce a high degree of stability for the values of the statistics of interest). This procedure was also used for the integrals involved in obtaining the load statistics from the probability distribution of allele frequencies. No integration over *γ* values is needed for Zone 1, since neutrality is assumed and the integral of *γ* over this interval is obtained using Equation (9).

### 2. Load calculations for a subdivided population

The load calculations for a subdivided population are divided into four different zones according to the scaled strength of selection. Zone 1 is such that the allele frequencies within both *In* and *St* over the metapopulation are effectively neutral. This case requires 0 ≤ 2*N*_*T*_*ys ≤ γ*_*c*1_, where *γ*_*c*1_ is the assigned threshold scaled selection coefficient for the metapopulation that defines effective neutrality, generally set to 0.25. The general formulae for the load statistics given by Equations (1) imply that, for a given *s* and *h*, they can be found from the expectations over the probability distributions of 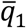 and 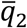, the mean frequencies of *A*_2_ within *In* and *St* over the metapopulation and the *F*_*STi*_, the *F*-statistics for the variances of the allele frequencies among demes within karyotypes. Under neutrality, the expected value of 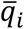 over the distribution of mean allele frequencies is 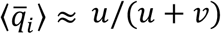 for both *In* and *St*, and allele frequencies within demes follow a beta distribution with this expectation (Wakeley 2003).

The *F*_*STi*_ can be calculated from the neutral formula for a diallelic locus:

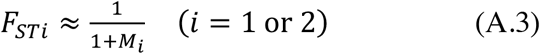

where *M*_1_ = 4*Nmx, M*_2_ = 4*Nmy*.

The net *F*-statistics for use in Equations (1) are then given by the standard result for hierarchical *F*-statistics (Wright 1951):

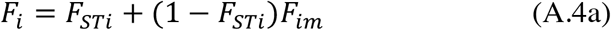

where *F*_*im*_ is the ratio of the variance in the 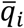 to the product of the expectations of 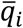 and 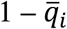. In the case of neutrality, we have:

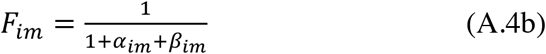

The expected load statistics contributed by zone 1 are then obtained as described for the case of a single population. By Maruyama’s invariance principle for neutral sites (Maruyama 1971; Charlesworth and Charlesworth 2010, p.318), the expected neutral diversity within a deme and karyotype is given by Equations (11), with *N*_*T*1_ and *N*_*T*1_ replacing *N*_1_ and *N*_2_, provided that *α*_*im*_ = 4*N*_*Ti*_ *u/*(1 − *F*_*STi*_) and *β*_*im*_ = 4*N*_*Ti*_ *v/*(1 − *F*_*STi*_) are both << 1. In other cases, the more general expression 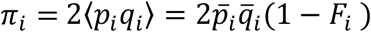 is to be used. As explained in the main text, the neutral values for the *F*_*STi*_ are also used in the calculations involving selection.

For Zone 2, the allele frequencies for both *In* and *St* within demes are effectively neutral, whereas the mean allele frequencies for the metapopulation are affected by moderate selection. The selection coefficients for zone 2 are now such that *γ*_*c*1_ *≤* 2*N*_*T*_*sy* ≤ *γ*_*c*2_, where *γ*_*c*2_ = *dγ*_*c*1_. The bivariate p.d.f. for 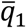 and 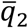, described by Equations (13), together with the relevant mutational terms, is used to determine their means and variances (see the Supplementary Information, section 1 for further details). For values of 2*N*_*T*_*sy* close to the lower boundary, the approximation described for zone 2a of the single population case was used. Since the distributions of allele frequencies within demes are close to neutrality, the covariance between 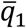 and 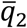 can be ignored. The variance of 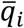 normalised by 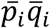 yields *F*_*im*_, the *F*-statistic for the metapopulation distribution. This can then be combined with the neutral value of *F*_*STi*_ from the beta distribution, and the expected load statistics obtained as described for zone 1.

Zone 3 involves moderate selection, where coupling between *In* and *St* in the metapopulation is ignored, but is allowed for the within-deme allele frequencies. In this case, the upper limit to *γy* is *γ*_*c*3_ = *γ*_*c*2_ /*x*. 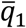 obeys Equations (13) with *a*_1_ = [*G*_1_ + (1 − 2*G*_1_)*h*]*x*^2^, 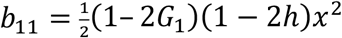 and the other terms are set to zero, so that the distributions of 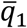 and 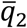 are treated as independent; this is justified by the fact the covariance *C*_12_ is always very small compared to the variances. Similarly, 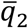 obeys Equations (13) with *a*_2_ = [*G*_2_ + (1 − 2*G*_2_)*h*]*y*^2^, 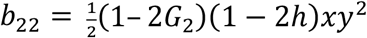.

In order to calculate the unconditional expected load statistics, the following approximation avoids integrations over the entire range of the 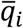. The distribution of 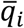 is divided into three regions, two of which are boundary regions with 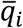 close to 0 or 1, respectively, and an intermediate region, denoted by labels 1, 3 and 2, respectively. We then use the expected values of 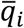 for each region to determine the corresponding values of the mutation-migration parameters *α*_*id*_ and *β*_*i*_ associated with Equations (13). These parameters are then used to obtain the values of the means, variances and covariance of the *q*_*i*_ across demes, integrating over the bivariate p.d.f. of Equations (13) as described in the Supplementary Information, section 1. The results are then substituted into Equation (1) to yield the load statistics for a given *s*. The means of these over the three regions are obtained by weighting by the probabilities of the 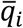 falling into the respective regions, and integration of over zone 3 by the same method as before.

The probabilities of the three regions and the corresponding expectations of the 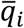 can be found as follows, noting that the assumption of independence between the two distributions means that the probability that 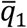 falls in region *i* and 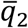 into region *j* is simply the product of the probabilities for each 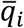 taken separately. In order to avoid singularities for values of 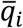 close to 0 or 1, it is convenient to approximate the p.d.f. for 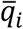 as follows:

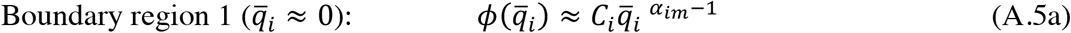

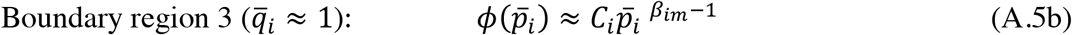

where *C*_*i*_ is the normalization constant for the p.d.f. of 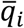. The *C*_*i*_ are obtained using a procedure like that described for the bivariate distribution in the Supplementary Information.

The probability that 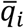 falls below an assigned critical value 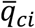, and hence into boundary region 1, is given by:

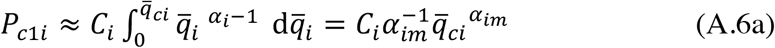

The expectation of 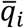 in this region is given by:

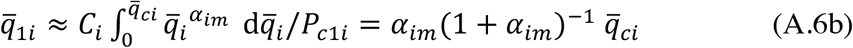

Similarly, the probability that 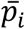 falls below an assigned critical value 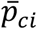, and hence into boundary region 3, is given by:

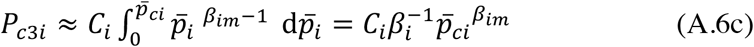

The expectation of 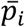 in this region is given by:

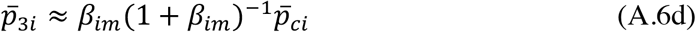

The probability that 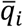 falls into the intermediate region 2 is *P*_2*i*_ = 1 − *P*_*c*1*i*_ − *P*_*c*3*i*_. The overall expectation of 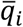, 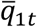, is obtained by integrating the modified version of Equations (13), similar to the procedure described for the bivariate distribution in the Supplementary Information. The expectation for the intermediate region 2 is then given by:

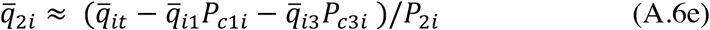

In practice, the critical values of all four mean allele frequencies are set to 0.001 or 0.01/*γ*, whichever is smaller, providing a conservative cut-off for neutrality.

For zone 4, both the *In* and *St* mean allele frequencies over the metapopulation are subject to strong selection. The lower bound to this region is *γ*_c3_ = *dγ*_c2_ /*x*. The upper bound *γ*_c4_ is chosen to correspond to the 99^th^ percentile of the distribution of *γ*. Both the *In* and *St* metapopulation distributions are assumed to be gamma distributions with means equal to *u*/*hs* and shape parameters *α*_1*m*_ and *α*_*2m*_, respectively Otherwise, the treatment of the distributions for the metapopulation, and the within-deme distributions are the same as for zone 3. The standard procedure for evaluating the total contribution to the load statistics is used.

